# Divergent pallidal pathways underlying distinct Parkinsonian behavioral deficits

**DOI:** 10.1101/2020.11.27.401554

**Authors:** Varoth Lilascharoen, Eric Hou-Jen Wang, Nam Do, Stefan Carl Pate, Amanda Ngoc Tran, Xiao-Yun Wang, Young-Gyun Park, Kwanghun Chung, Byung Kook Lim

## Abstract

The basal ganglia are a group of subcortical nuclei that regulates motor and cognitive functions^1,2^. Recent identification of neuronal heterogeneity in the basal ganglia suggests that functionally distinct neural circuits defined by their efferent projections exist even within the same nuclei^3-5^. This distinction may account for a multitude of symptoms associated with basal ganglia disorders such as Parkinson’s disease (PD)^6,7^. However, our incomplete understanding of the basal ganglia functional organization has hindered further investigation of individual circuits that may underlie different behavioral symptoms in disease states. Here we functionally define two distinct classes of parvalbumin-expressing neurons in the mouse external globus pallidus (GPe-PV) embedded within discrete neural pathways and establish their contributions to different Parkinsonian behavioral deficits. We find that GPe-PV neurons projecting to the substantia nigra pars reticulata (SNr) or parafascicular thalamus (PF) undergo different electrophysiological adaptations in response to dopamine depletion. Furthermore, counteracting these adaptations in each population can selectively alleviate movement deficits or behavioral inflexibility in a Parkinsonian mouse model. Our findings provide a novel framework to understand the circuit basis of separate behavioral symptoms in Parkinsonian state which could provide better strategies for the treatment of PD.

---

The GPe is a central basal ganglia nucleus that can influence numerous downstream regions^8^. The majority of GPe neurons express parvalbumin and innervate multiple basal ganglia and thalamic nuclei^9-12^. Recent studies have demonstrated that projection-defined neuronal subclasses within the neighboring internal globus pallidus (GPi) and ventral pallidum can mediate separate behaviors^3-5^, suggesting that a similar functional organization may exist within the GPe. However, the relationship between the anatomical organization of GPe neurons and behavioral function remains poorly understood.

Great attention has been drawn to the potential role of the GPe in motor control as movement-entrained neural activity have been observed in GPe neurons. Moreover, aberrant GPe neuronal activity is highly associated with movement deficits in both PD patients^13^ and Parkinsonian animal models^14,15^. Importantly, high-frequency deep-brain stimulation of the GPe in human patients and prolonged optogenetic activation of the GPe-PV neurons in dopamine-depleted mice can alleviate Parkinsonian motor symptoms^16,17^. These stimulation effects may be mediated through GPe neurons that innervates the substantia nigra pars reticulata (SNr), an output nucleus of the basal ganglia that directly controls motor-related brain regions, including the motor thalamus and the mesencephalic locomotor region^18,19^. Moreover, a previous study demonstrated that a subset of GPe neurons displayed a robust activity dynamic during a task that requires stimulus-action-outcome association^20^. This raises the intriguing possibility that control of non-motor behavior may also be achieved through GPe-PV neurons that innervate non-motor related nuclei such as the parafascicular thalamus (PF), a nucleus known to be involved in cognitive function^21,22^. Together, this suggests that GPe-PV neurons may influence a variety of behavioral functions through distinct neuronal populations, yet our incomplete understanding of the GPe circuit organization has precluded any investigation of a potential divergent control of motor and non-motor behaviors. Here, we investigate the projection-specific roles of the GPe-PV neurons and their contributions to motor and behavioral flexibility impairments in a Parkinsonian mouse model.

## Divergent pathways of GPe-PV neurons

We first examined the projection pattern and synaptic targets of GPe-PV neurons by injecting an adeno-associated virus (AAV) expressing a red fluorescent protein, mRuby2, and synaptophysin fused with eGFP in a Cre-dependent manner into the GPe of *PV*^*Cre*^ mice. This approach restricts eGFP expression to putative presynaptic terminals, allowing us to discriminate between actual synaptic targets and axon-of-passage^4^. Consistent with previous findings, we observed robust labeling of synaptic terminals in the subthalamic nucleus, GPi, SNr, PF, and dorsal striatum (Fig. 1a-b)^12,23^.

**Fig. 1.**
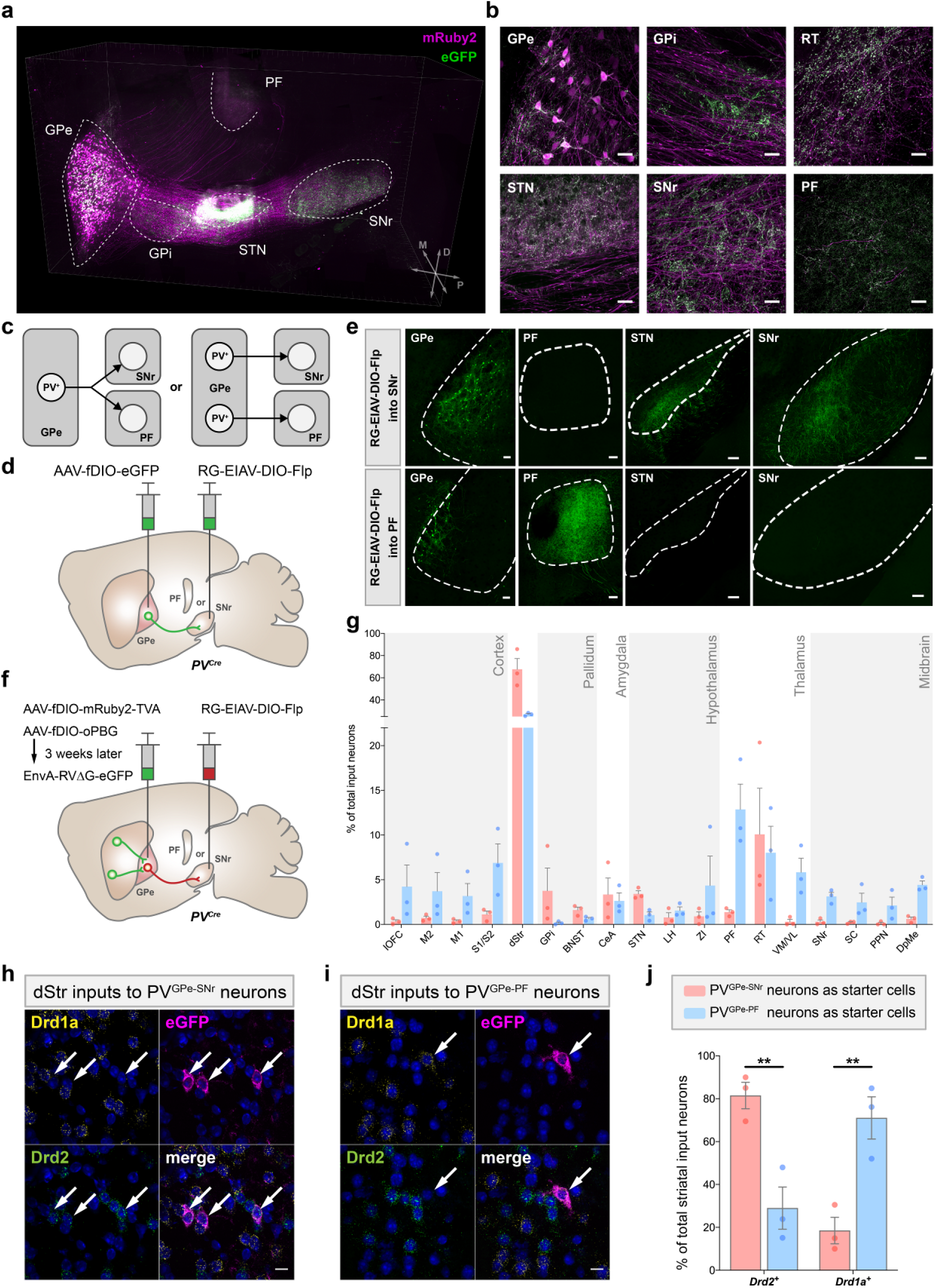
Distinct subpopulations of GPe-PV neurons project to the SNr and PF. **a**, 3D rendering of a cleared mouse hemisphere showing brain-wide projection patterns of GPe-PV neurons labeled by mRuby2 (soma, axonal fibers) and eGFP (presynaptic sites). **b**, Representative confocal images of the injection site in GPe (top left) and target structures showing axonal fibers in magenta and synaptic puncta in green. Scale bar, 30 μm. **c**, Two possible projection patterns: GPe PV neurons collateralizes to multiple targets (left), or individual neurons project to distinct targets (right). **d**, Schematic of viral strategy using a pseudotyped equine infectious anemia lentivirus capable of neuron-specific retrograde infection (RG-EIAV). SNr or PF of *PV*^*Cre*^ mice is injected with RG-EIAV that expresses Flp recombinase in a Cre-dependent manner (RG-EIAV-DIO-Flp) and the GPe with an AAV that expresses eGFP in a Flp-dependent manner (AAV-fDIO-eGFP). **e**, Confocal images of SNr- and PF-projecting GPe-PV neurons and their axons. Scale bar, 100 μm. **f**, Viral strategy to map pseudotyped rabies-mediated monosynaptic inputs to PV^GPe-SNr^ neurons. In *PV*^*Cre*^ mice, SNr or PF was injected with RG-EIAV-DIO-Flp and GPe was injected with AAV-fDIO-TVA-mRuby, AAV-fDIO-oPBG. EnvA-RVΔG-eGFP was subsequently injected into the GPe. **g**, Whole-brain quantification of inputs to PV^GPe-SNr^ and PV^GPe-PF^ neurons. Data presented as percentage of total cells in each brain area relative to the total number of brain-wide inputs (*n* = 3 mice for each subpopulation). **h-i**, Representative images of striatal neurons sending input to PV^GPe-SNr^ (**h**) and PV^GPe-PF^ (**i**) neurons with cell-type-specific markers. Arrows represent colocalization between striatal input neurons labeled by the rabies virus and mRNA labeled by cell-type-specific probes. **j**, Quantification of *Drd1a*^*+*^ or *Drd2*^*+*^ striatal inputs to PV^GPe-SNr^ and PV^GPe-PF^ neurons. Paired *t*-test, t(2) = 9.333; \**p* = 0.0113 (*n* = 3 mice for each subpopulation). Scale bars, 30 μm (**b**), 100 μm (**e**), and 10 μm (**h-i**). All data presented as mean ± SEM. lOFC, lateral orbitofrontal cortex; M2, secondary motor cortex; M1, primary motor cortex; S1/S2, somatosensory cortex; dStr, dorsal striatum; GPi, internal globus pallidus; BNST, bed nucleus of stria terminalis; CeA, central amygdala; STN, subthalamic nucleus; LH, lateral hypothalamus; ZI, zona incerta; RT, reticular thalamus; VM/VL, ventromedial/ventrolateral thalamus; SC, superior colliculus; PPN, pedunculopontine nucleus; DpMe, deep mesencephalic nucleus.

Although GPe neurons project to many regions, it remains unclear whether separate GPe neurons project onto distinct targets or individual GPe neurons collateralize onto multiple targets (Fig. 1c). In particular, we focused on the GPe outputs to the SNr and PF, two regions known to be involved in motor control^18^ and behavioral flexibility^21,22^, respectively. These distinct functions led us to hypothesize that distinct subclasses of GPe-PV neurons project to the SNr or PF. To test this hypothesis, we used an intersectional genetic targeting approach to label GPe neurons in a cell type- and projection-specific manner (Fig. 1d)^4,24,25^. By restricting eGFP expression to a projection-defined subset of GPe-PV neurons, we found that SNr-projecting GPe-PV neurons (PV^GPe-SNr^) were present throughout the GPe and innervate multiple nuclei in the basal ganglia: STN and GPi. By contrast, the PF-projecting GPe-PV neurons (PV^GPe-PF^) were clustered in the ventromedial region of the GPe and innervate only the PF (Fig. 1e). Thus, PV^GPe-SNr^ and PV^GPe-PF^ neurons are distinct neuronal populations that may represent separate neural pathways.

## Inputs to PV^GPe-SNr^ and PV^GPe-PF^ neurons

To determine whether PV^GPe-SNr^ and PV^GPe-PF^ neurons are, indeed, part of separate neural pathways, we aimed to demonstrate that each population receives inputs from different brain regions. We mapped the brain-wide inputs to each subpopulation by expressing the TVA receptor and an optimized rabies glycoprotein (oPBG)^26^ in either PV^GPe-SNr^ or PV^GPe-PF^ neurons followed by an injection of the EnvA-pseudotyped, glycoprotein-deleted rabies virus (EnvA-RVΔG-eGFP) into the GPe (Fig. 1f). Whole-brain quantification of eGFP-labeled neurons revealed that the PV^GPe-SNr^ neurons receive proportionally more inputs from the STN, whereas the PV^GPe-PF^ neurons receive more inputs from the cortex and the midbrain (Fig. 1g, Extended Data Fig. 1). While the main input of both populations originates in the striatum, PV^GPe-SNr^ and PV^GPe-PF^ neurons are preferentially innervated by the dorsolateral and dorsomedial striatum, respectively (Extended Data Fig. 1). Furthermore, by determining the molecular identity of the striatal input neurons, we found that PV^GPe-SNr^ and PV^GPe-PF^ neurons receive proportionally more inputs from striatal neurons expressing *Drd1a* or *Drd2a* mRNA, respectively. Taken together, this suggests that PV^GPe-SNr^ and PV^GPe-PF^ neurons are part of distinct neural pathways that may underlie different behavioral functions. (Fig. 1h-j).

## Distinct properties of PV^GPe-SNr^ and PV^GPe-PF^ neurons

GPe neurons exhibit heterogeneous autonomous firing patterns and intrinsic membrane properties^10,11,27^. To determine whether PV^GPe-SNr^ and PV^GPe-PF^ neurons also differ in their electrophysiological properties, we performed *ex vivo* whole-cell recordings in acute brain slices. We selectively labeled PV^GPe-SNr^ and PV^GPe-PF^ neurons by injecting RG-EIAV expressing Cre in a Flp-dependent manner (RG-EIAV-fDIO-Cre) into either SNr or PF of *PV*^*Flp*^ *x Ai14* mice, resulting in projection-specific tdTomato expression in GPe-PV neurons (Extended Data Fig. 2a). Both populations were spontaneously active at rest with the PV^GPe-SNr^ neurons firing more regularly at a higher rate compared to the PV^GPe-PF^ neurons (Extended Data Fig. 2b-d). Further evaluation also revealed differences in driven activity and action potential waveforms of these two subpopulations (Extended Data Fig. 2e-l). Thus, in addition to their different anatomical connectivity, PV^GPe-SNr^ and PV^GPe-PF^ neurons are distinguishable by their electrophysiological properties.

## PV^GPe-SNr^ neurons mediate locomotion

Given their distinct connectivity patterns and electrophysiological properties, PV^GPe-SNr^ and PV^GPe-PF^ neurons may convey separate information streams to their respective targets to mediate different behaviors. We aimed to determine whether the activity of PV^GPe-SNr^ and PV^GPe-PF^ neurons displayed a time-locked response to locomotion initiation and termination. We expressed axon-targeted, genetically encoded Ca^2+^ indicator (axon-GCaMP6s)^28^ and mRuby3 in GPe-PV neurons and used fiber photometry to simultaneously measure the population calcium activity from their axon terminals in the SNr and PF during locomotion (Fig. 2a, Extended Data Fig. 3a-b)^29,30^. Mice were head-fixed over a cylindrical treadmill, and GCaMP6s fluorescence was measured during self-initiated locomotion (Fig. 2b). We observed an increase in fluorescence intensity in both target areas during the transition from rest to run. Although the activity of both subpopulations is correlated with locomotion bouts, the rise in PV^GPe-SNr^ neuronal activity occurred slightly before the locomotion onset, while the PV^GPe-PF^ neuronal activity rose after the onset (Fig. 2c-d). This suggests that both populations may differentially mediate locomotion.

**Fig. 2.**
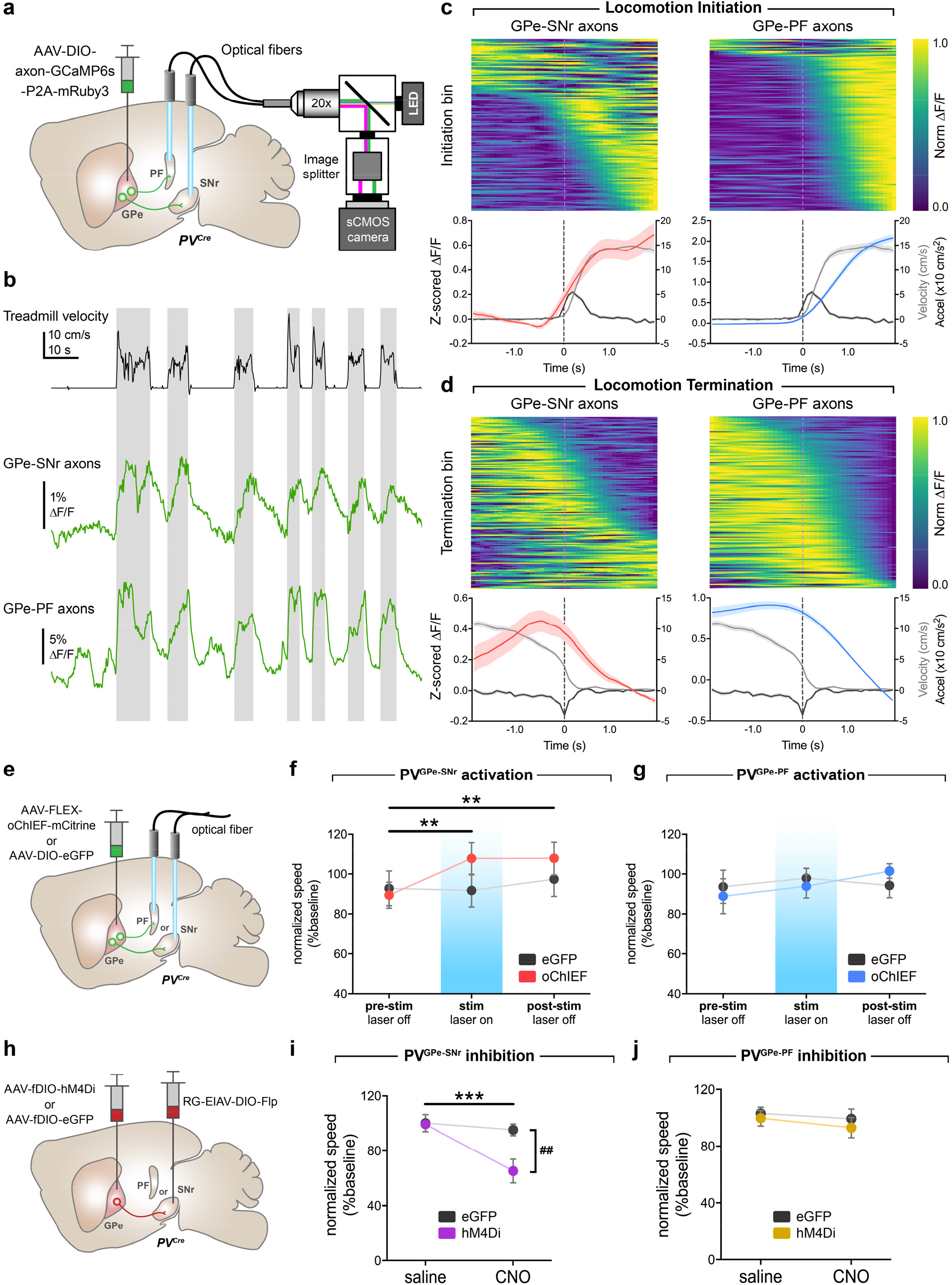
Activity of PV^GPe-SNr^ neurons bidirectionally modulates locomotion. **a**, Schematic of fiber photometry setup. **b**, Representative ΔF/F traces of GCaMP6s fluorescence recorded from axonal fibers of PV^GPe-SNr^ and PV^GPe-PF^ neurons from one mouse (*n* = 7 mice total) and corresponding treadmill velocity (top). Shaded areas indicated locomotion bins. **c**, Top, ΔF/F aligned to locomotion initiations from all mice (ΔF/F was normalized for each row and sorted by time to half-max). Bottom, mean acceleration (black), velocity (grey), and z-scored ΔF/F triggered at locomotion initiation (mean across all events). **d**, same as **c**, except aligned to locomotion terminations. **e**, Schematic of viral injections and optic fiber implantations for optogenetic activation. **f**, Activation of GPe PV terminal in the SNr increased locomotion during stimulation and post-stimulation periods. Two-way repeated-measures ANOVA, main effect_stimxgroup_: *F*(2,36) = 3.863, *p* = 0.0302; main effect_stim_: *F*(2,36) = 5.513, *p* = 0.0082; main effect_group_: *F*(1,18) = 0.5161, *p* = 0.4817; Bonferroni’s *post hoc* test, \*\**p* < 0.01; *n* = 10 mice for both eGFP and oChIEF). **g**, Activation of GPe PV terminal in the PF had no effect on locomotion. Two-way repeated-measures ANOVA, main effect_stimxgroup_: *F*(2,26) = 0.6664, *p* = 0.5221; main effect_stim_: *F*(2,26) = 0.6870, *p* = 0.5120; main effect_group_: *F*(1,13) = 0.0065, *p* = 0.9368; *n* = 7 mice for eGFP and *n* = 8 mice for oChIEF). **h**, Schematic of viral injections for cell-type- and projection-specific expression of inhibitory DREADD (hM4Di) in PV^GPe-SNr^ and PV^GPe-PF^ neurons. **i**, Inhibition of PV^GPe-SNr^ neurons suppresses locomotion. Two-way repeated-measures ANOVA, main effect_treatmentxgroup_: *F*(1,18) = 9.644, *p* = 0.0061; main effect_treatment_: *F*(1,18) = 17.30, *p* = 0.0006; main effect_group_: *F*(1,18) = 4.968, *p* = 0.0388; Bonferroni’s *post hoc* test, \*\*\**p* < 0.001; ^##^*p* = 0.002; *n* = 10 mice for both eGFP and hM4Di). **j**, Inhibition of PV^GPe-PF^ neurons had no effect on locomotion. Two-way repeated-measures ANOVA, main effect_treatmentxgroup_: *F*(1,10) = 0.069, *p* = 0.7980; main effect_treatment_: *F*(1,10) = 0.9681, *p* = 0.3484; main effect_group_: *F*(1,10) = 0.5119, *p* = 0.4911; *n* = 6 mice for both eGFP and hM4Di). Data in **f-j** are presented as mean speed (normalized to baseline) ± SEM. Shaded areas in **c-d** indicate SEM.

To unambiguously determine whether the activity of PV^GPe-SNr^ and PV^GPe-PF^ neurons can modulate locomotion, we photostimulated oChIEF-expressing PV^GPe^ axons in the SNr or PF and observed significantly increased locomotor activity during and after SNr but not PF stimulation (Fig. 2e-g). Next, we tested whether the activity of PV^GPe-SNr^ and PV^GPe-PF^ neurons is necessary for locomotion. We utilized chemogenetic inhibition by expressing the inhibitory designer receptor (hM4Di) in the PV^GPe-SNr^ or PV^GPe-PF^ neurons that can be activated by clozapine *N*-oxide (CNO; Fig. 2h). Inactivation of the PV^GPe-SNr^ neurons induced a significant reduction in locomotor activity, while inactivation of the PV^GPe-PF^ neurons did not affect locomotion (Fig. 2i-j). Collectively, our results indicated that the activity of PV^GPe-SNr^ neurons, but not PV^GPe-PF^ neurons, can bidirectionally influence locomotion and plays an essential role in motor control.

## PV^GPe-PF^ neurons are involved in behavioral flexibility

To investigate the involvement of GPe-PV neurons in behavioral flexibility, we used fiber photometry to examine the activity of both subpopulations during a naturalistic foraging reversal discrimination task. Mice were first trained to associate a certain context with a food reward, and a different context without reward until they can discriminate reliably (association phase). The behavioral flexibility was determined in the subsequent reversal-learning phase in which the context-outcome associations were reversed (Fig. 3a, Extended Data Fig. 4a-b)^31^. The overall activity of PV^GPe-PF^ neurons was significantly higher during trials relative to the rest periods between trials (inter-trial intervals; ITIs), while there was no difference in the activity of the PV^GPe-SNr^ neurons between the two periods. This suggests that the activity of PV^GPe-PF^ neurons is selectively associated with the task (Extended Data Fig. 4c-d). Next, we examined the time course of the activity event-locked to trial start or the moment that the mice started digging to signify a choice (dig start; Fig. 3b) in three different stages of the task—association, early reversal learning (a period before the criterion was met), and late reversal learning (a period at which the criterion was met; details in Methods). Both subpopulations showed a brief event-locked response to trial onsets across all stages of the task, possibly due to an increase in locomotion at the start of each trial (Fig. 3c, e). Notably, we found that only the PV^GPe-PF^ neurons showed a significant increase in activity time-locked to when the mice started digging in the early stage of reversal learning relative to the association and late stage of reversal learning (Fig. 3d, f).

**Fig. 3.**
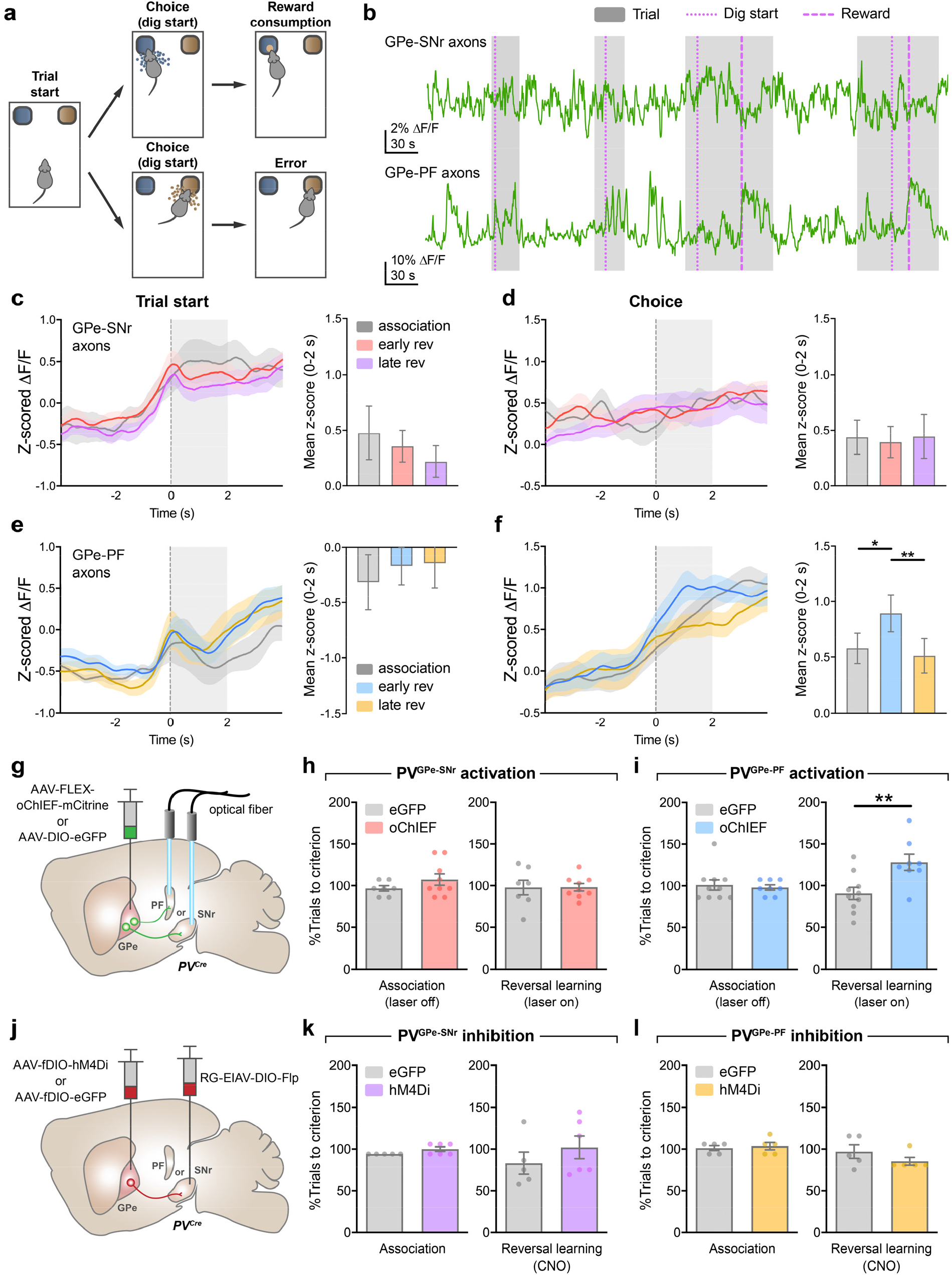
Activation of PV^GPe-PF^ neurons impairs behavioral flexibility in reversal learning. **a**, Behavioral task schematic showing the structure of each trial; after trial starts, the mouse makes a choice by digging in one of the two bowls. Only correct choice yields a food reward. **b**, Representative fiber photometry ΔF/F traces showing the activity of PV^GPe-SNr^ and PV^GPe-PF^ axons from one mouse (*n* = 7 mice total) encompassing 4 trials in the reversal-learning phase. Thin dotted line, thick dotted line, and shaded area represent dig start, reward consumption, and trial period, respectively. **c-d**, Left, traces of z-scored ΔF/F (averaged across 7 mice) from PV^GPe-SNr^ axons aligned to trial start (**c**) and timing of choice (**d**). Right, mean z-score from 0-2 s after behavioral onset. Early rev, early stage of reversal learning; Late rev, late stage of reversal learning. One-way repeated-measures ANOVA, effect_stage_: *F*(2,12) = 1.300, *p* = 0.3084 in **c**, and effect_stage_: *F*(2,12) = 0.0378, *p* = 0.9630 in **d. e-f**, Activity of PV^GPe-PF^ axons aligned to trial start (**e**) and timing of choice (**f**) same as in **c-d**. One-way repeated-measures ANOVA, effect_stage_: *F*(2,12) = 1.280, *p* = 0.3135 in **e**, and effect_stage_: *F*(2,12) = 8.008, *p* = 0.0062 in **f**; Bonferroni’s *post hoc* test, \**p* = 0.0279, \*\**p* = 0.0084. **g**, Schematic of viral injections and optic fiber implantations for optogenetic activation during reversal learning. **h**, Activation of PV^GPe-SNr^ axons did not affect behavioral flexibility during reversal learning. Unpaired *t*-test, *t*(16) = 0.0273, *p* = 0.9786; *n* = 7 mice for eGFP and *n* = 9 mice for oChIEF. **i**, Activation of PV^GPe-PF^ axons during reversal learning impaired behavioral flexibility. Unpaired *t*-test, *t*(16) = 3.142, \*\**p* = 0.0063; *n* = 10 mice for eGFP and *n* = 8 mice for oChIEF. **j**, Schematic of viral injections and experimental timeline for cell-type- and projection-specific expression of inhibitory DREADD (hM4Di) in PV^GPe-SNr^ and PV^GPe-PF^ neurons. **k**, Inhibition of PV^GPe-SNr^ neurons during reversal learning had no effect on behavioral flexibility. Unpaired *t*-test, *t*(9) = 0.9840, *p* = 0.3508; *n* = 5 mice for eGFP and *n* = 6 mice for hM4Di. **l**, Inhibition of PV^GPe-PF^ neurons during reversal learning had no effect on behavioral flexibility. Unpaired *t*-test, *t*(8) = 1.245, *p* = 0.2483; *n* = 5 mice for both eGFP and hM4Di. Shaded areas accompanying the z-scored ΔF/F traces in **c-f** indicate SEM. All other data are presented as mean ± SEM.

The changes in activity of PV^GPe-PF^ neurons when the mice signified a choice across different stages of the task raised the question of whether the activity dynamic of these neurons is involved in mediating behavioral flexibility. To answer this question, we manipulated the activity dynamics during reversal learning by prolonged optogenetic or chemogenetic manipulation and assess the behavioral consequences. While photostimulation of PV^GPe-SNr^ axons did not affect behavioral flexibility, activation PV^GPe-PF^ axons significantly increased the number of trials to criterion (Fig. 3g-i). Further analysis revealed a higher number of regressive errors when the PV^GPe-PF^ axons were activated, indicating that the mice had difficulty in maintaining the new context-outcome association (Extended Data Fig. 6). Chemogenetic inhibition of both PV^GPe-SNr^ and PV^GPe-PF^ neurons during reversal learning did not affect behavioral flexibility (Fig. 3j-l). Our results indicated that, although not required during reversal learning, sustained activation of PV^GPe-PF^ neurons can negatively affect behavioral flexibility.

## Dopamine depletion alters synaptic output of GPe-PV neurons

Patients with PD exhibit a multitude of behavioral symptoms that likely reflect the adaptations in different neural circuitries. The differential effects of projection-specific manipulation of GPe-PV neurons on locomotion and behavioral flexibility suggest that alteration in synaptic inputs, outputs, or intrinsic properties of PV^GPe-SNr^ and PV^GPe-PF^ neurons following dopamine depletion may underlie different parkinsonian behavioral deficits^32,33^. To induce dopamine depletion, we used a well-established 6-hydroxydopamine-(6-OHDA)-lesioned Parkinsonian mouse model. We first recorded from the PV^GPe-SNr^ and PV^GPe-PF^ neurons in acute brain slices and found that dopamine depletion had no effect on autonomous firing rate and the ratio of excitatory and inhibitory synaptic inputs (E/I ratio) of the PV^GPe-SNr^ and PV^GPe-PF^ neurons (Extended Data Fig. 8a-e). Thus, we subsequently examined whether synaptic outputs at the GPe-SNr and GPe-PF synapses were altered. We replaced extracellular Ca^2+^ with Sr^2+^ and photostimulated GPe-PV axons while recording from SNr and PF neurons to measure quantal-like inhibitory postsynaptic currents (qIPSC) from the stimulated synapses^34^. Interestingly, we observed a significant decrease in the frequency of qIPSC in SNr neurons, while that in the PF neurons is markedly increased (Fig. 4a-d). Additionally, the paired-pulse ratios of IPSCs, measured while photostimulating the GPe-PV axons, were increased in SNr neurons and decreased in PF neurons (Extended Data Fig. 8e-g). Taken together, our results indicated that dopamine depletion affected the synaptic outputs of PV^GPe-SNr^ and PV^GPe-PF^ neurons by altering the probability of presynaptic neurotransmitter release.

**Fig. 4.**
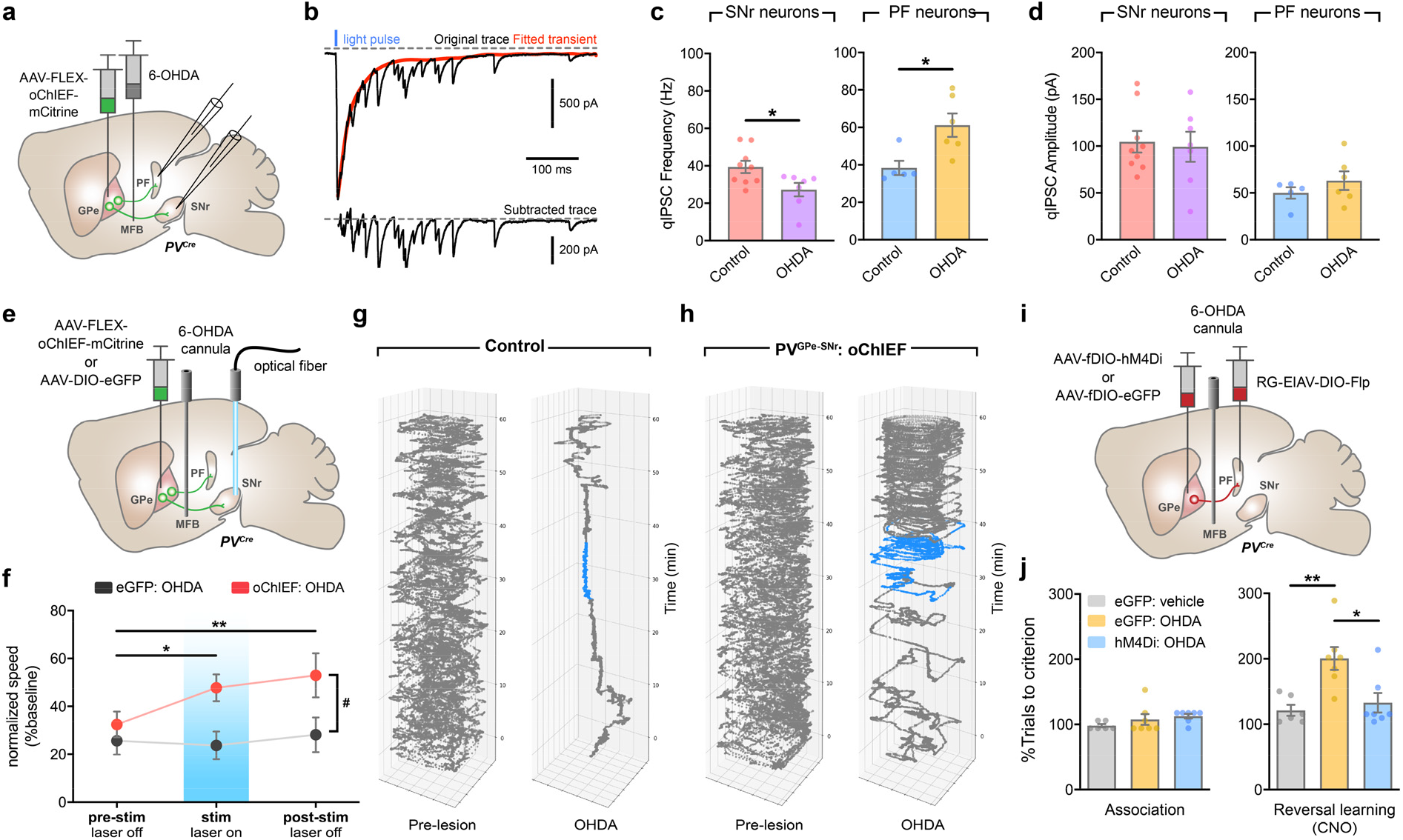
PV^GPe-SNr^ and PV^GPe-PF^ neurons exhibit distinct electrophysiological adaptations and mediate different behavioral deficits in dopamine depleted mice. **a**, Schematic of viral and 6-OHDA injections for measuring quantal-like inhibitory postsynaptic currents (qIPSC) in acute slices after dopamine depletion while photostimulating PV^GPe-SNr^ and PV^GPe-PF^ axons. **b**, Example trace (upper) showing quantal-like inhibitory postsynaptic currents (qIPSC) in SNr neuron elicited by photostimulation of GPe-PV terminals. Red trace represents a fitted curve generated by Python script. qIPSC amplitudes were measured from subtracted trace (lower) from 0-400 ms after stimulation. **c**, Frequency of optically-evoked qIPSC was decreased and increased in when recorded from SNr and PF neurons, respectively. Unpaired *t*-test, *t*(14) = 2.485, \**p* = 0.0262 for SNr, and *t*(9) = 2.947, \**p* = 0.0163 for PF (SNr control *n* = 9 cells from 3 mice, SNr OHDA *n* = 7 cells from 3 mice; PF control *n* = 5 cells from 2 mice, PF OHDA *n* = 6 cells from 2 mice). **d**, Amplitudes of optically-evoked qIPSC in SNr and PF neurons were not altered by dopamine depletion. Unpaired *t*-test, *t*(14) = 0.2773, *p* = 0.7856 for SNr, and *t*(9) = 1.052, \**p* = 0.3204 for PF (SNr control *n* = 9 cells, SNr OHDA *n* = 7 cells; PF control *n* = 5 cells, PF OHDA *n* = 6 cells). **e**, Schematic of viral injections, optic fiber, and drug cannula implantations for optogenetic activation during locomotion in dopamine-depleted mice. **f**, Activation of PV^GPe-SNr^ axons restored locomotor activity during stimulation and post-stimulation periods in dopamine-depleted mice. Two-way repeated-measures ANOVA, main effect_stimxgroup_: *F*(2,38) = 2.691, *p* = 0.0807; main effect_stim_: *F*(2,38) = 3.460, *p* = 0.0417; main effect_group_: *F*(1,19) = 5.436, *p* = 0.0309; Bonferroni’s *post hoc* test, ^#^*p* < 0.05, \**p* < 0.05, \*\**p* < 0.01; eGFP *n* = 10 mice, oChIEF *n* = 11 mice). **g**, Position of a control mouse in an open field over 60 min before (left) and after (right) dopamine depletion. The mouse received bilateral photostimulation in SNr at 30-40 min after session start (blue line). **h**, same as in **g**, except for a mouse with oChIEF expression in GPe-PV neurons. **i**, Schematic of viral injections and drug cannula implantations, for cell-type- and projection-specific expression of inhibitory DREADD (hM4Di) in PV^GPe-PF^ neurons. **j**, Performance of mice that received chemogenetic inhibition in PV^GPe-PF^ neurons after dopamine depletion. Mice in all groups received CNO injections (5 mg/kg) in the reversal-learning phase (*n* = 7 mice for eGFP-vehicle, *n* = 7 mice for eGFP-OHDA, and *n* = 7 mice for hM4Di-OHDA). Left, dopamine depletion did not affect performance in the association phase. Right, inhibition of PV^GPe-PF^ neurons during reversal learning improved behavioral flexibility in dopamine-depleted mice. One-way ANOVA, *F*(2,17) = 8.642, *p* = 0.0026; Bonferroni’s *post hoc* test, \**p* = 0.0114, \*\**p* = 0.0046. All data presented as mean ± SEM.

## Roles of GPe-PV neurons in parkinsonian behavioral deficits

The different behavioral contribution of PV^GPe-SNr^ and PV^GPe-PF^ neurons and their synaptic changes by dopamine depletion led us to speculate that these circuits underlie different Parkinsonian deficits. Our results strongly suggest that reduced synaptic output of PV^GPe-SNr^ neurons may lead to decreased locomotor activity. Thus, artificially enhancing the activity of PV^GPe-SNr^ neurons in dopamine-depleted mice may reverse locomotive deficits. In line with previously published works, we observed a significant reduction in locomotion when more than 70% of nigrostriatal dopaminergic fibers were lesioned (Fig. 4g-h and Extended Data Fig. 7)^35^. Interestingly, photostimulation of PV^GPe-SNr^ neurons in dopamine-depleted mice induced a robust increase in locomotion that persisted beyond the stimulation period (Fig. 4e-h), suggesting that reduced GPe-SNr synaptic output may indeed contribute to motor deficit in PD.

Besides motor deficits, PD patients and Parkinsonian animal models display cognitive flexibility impairments in the early stage of the disease^33,36,37^. Since PV^GPe-PF^ neurons showed increased synaptic output after dopamine depletion, inactivating these neurons may rescue behavioral inflexibility. To test this, we first induced a mild dopamine depletion by a low-dose 6-OHDA to mimic the early stage of dopamine loss (∼50% loss of striatal dopaminergic fibers; Extended Data Fig. 7). Mice did not display any locomotor deficit, thus allowing us to assess the behavioral flexibility with the reversal-learning task. We found that formation of the context-outcome association was not affected by dopamine depletion, whereas behavioral flexibility was profoundly impaired (Fig. 4j) as reflected by an increase in regressive errors (Extended Data Fig. 9a). We then chemogenetically inactivated PV^GPe-PF^ neurons which resulted in a significant improvement of behavioral flexibility following mild dopamine depletion (Fig. 4i-j, Extended Data Fig. 9a). By contrast, activation of the PV^GPe-SNr^ neurons showed no effect (Extended Data Fig. 9b-c). Collectively, these experiments revealed that each of the two subpopulations underwent specific adaptation in response to dopamine loss, which contributed to a separate behavioral deficit observed at a different stage of dopamine depletion.

## Concluding Remarks

Using complementary electrophysiological, viral-mediated tracing, and behavioral approaches, we have presented evidence that distinct circuits in the GPe are involved in different basal ganglia-related behaviors and that the physiological adaptation of each circuit contributes to a different behavioral deficit observed in a Parkinsonian mouse model. Our viral-mediated tracing indicated that GPe-PV neurons could be distinguished based on their projections to either SNr or PF. Interestingly, contrary to the classical model of the basal ganglia, we found that the PV^GPe-PF^ neurons are preferentially innervated by *Drd1a*-expressing striatal neurons^1^, suggesting that distinct GPe circuits can convey different information streams to their respective targets. Prior studies have focused mainly on how the basal ganglia output control the motor functions through thalamocortical loop and motor-related nuclei in the midbrain^38-40^. However, the functional roles of the direct output from the GPe to the thalamus is largely unknown. In addition to the involvement of the GPe-SNr pathway in locomotion, we find that the GPe-PF pathway is selectively involved in behavioral flexibility. This raises an intriguing possibility that, although the basal ganglia as a whole, is associated with a broad range of behaviors, functional roles of individual microcircuits within the basal ganglia can be different.

The progression of PD is commonly evaluated by the severity of motor symptoms; however, non-motor symptoms, such as cognitive impairments, are also common and often appear in an early phase of the disease^41^. Nevertheless, the potential involvement of different neural circuits in both motor or non-motor symptoms of PD is poorly understood. Here, we demonstrated that, GPe-SNr and GPe-PF synapses were differentially altered following dopamine depletion. By selectively counteracting these synaptic changes, we rescued locomotor deficit and behavioral inflexibility at different stages of Parkinsonian state, suggesting that the neural adaptations in different circuits may underlie the progressive nature of PD. Since our findings suggest that the GPe-PF pathway is involved in reversal learning, manipulating the activity of this pathway could provide a novel treatment for ameliorating behavioral inflexibility in PD. We believe that our findings establish the behavioral significance of two distinct GPe-PV neuronal populations embedded in discrete neural pathways and their differential contributions to specific subdomains of Parkinsonian-like behaviors that occur at different stages of the disease. This further suggests that evaluation of the detailed synaptic connectivity of the GPe is needed to fully understand the circuit-specific roles of the basal ganglia, and could provide better therapeutic strategies for the treatment of PD.

## Acknowledgments

We thank D. Knowland, C. Santiago, and I. Zutshi for their comments on the manuscript. S. Lilascharoen helped with quantification in Fig. 1 and illustrations. We thank the members of the Lim laboratory for support and discussions. V.L. was supported by the Anandamahidol Foundation Fellowship. This work was supported by grants from NIH (NS094342, NS097772 and MH114829).

## Author contributions

V.L. and B.K.L. conceived and designed the study. V.L. performed all experiments, analyzed the data, and interpreted the results. V.L., E.H.W., S.C.P., and A.N.T performed stereotaxic surgery. V.L. and E.H.W. performed behavioral experiments. V.L., S.C.P., A.N.T., and X.-Y.W designed and generated viruses. V.L., N.D., S.C.P., and A.N.T. performed histology and immunohistochemistry. E.H.W. constructed the fiber photometry recording system and assisted with data analysis. V.L., S.C.P., and Y.-G.P performed SHIED-MAP tissue clearing and light-sheet imaging. Y.-G.P and K.C. provided resources for SHIELD-MAP. V.L. and B.K.L. wrote the manuscript with contributions from E.H.W., S.C.P., and A.N.T.

## METHODS

### Animals

All procedures to maintain and use mice were approved by the Institutional Animal Care and Use Committee (IACUC) at the University of California, San Diego. Mice were maintained on a 12-hour light/dark cycle with regular mouse chow and water available *ad libitum* except when placed under food restriction. All behavioral experiments were performed during the dark cycle. *PV*^*Cre*^, *Ai14* (Rosa26-CAG-LSL-tdTomato), and *PV*^*Flp*^ transgenic mice were obtained from the Jackson Laboratory (JAX strain 008069, 007914, and 22730, respectively) and were maintained on a C57BL/6J background. *PV*^*Flp*^ *x Ai14* transgenic mice were generated by crossing *PV*^*Flp*^ mice with *Ai14* mice. For all experiments, male and female heterozygous mice aged 8-20 weeks were used.

### Viral vectors

AAV plasmids were constructed using standard molecular cloning methods. Synaptophysin-eGFP, oPBG, hM4D(Gi)-mCherry, axon-GCaMP6s-P2A-mRuby3, and TVA DNA fragments were obtained from pAAV-phSyn1(S)-FLEX-tdTomato-T2A-SypEGFP-WPRE (a gift from Hongkui Zeng; Addgene plasmid #51509), pAAV-hSyn-DIO-hM4D(Gi)-mCherry (a gift from Bryan Roth; Addgene plasmid #44362), pAAV-EF1α-DIO-oPBG (a gift from Edward M. Callaway), pAA-hSynapsin1-axon-GCaMP6s-P2A-mRuby3 (a gift from Lin Tian; Addgene plasmid #112005), and pAAV-EF1α-FLEX-GTB (a gift from Edward M. Callaway; Addgene plasmid #26197), respectively. pAAV-FLEX-oChIEF-mCitrine was a gift from Roger Tsien (Addgen plasmid #50973). A Flp-dependent, double-floxed, inverted open reading frame (fDIO) was constructed with two heterospecific pairs of FRT and FRT5 sequences based on pAAV-EF1α-fDIO-hChR2(H134R)-eYFP (a gift from Karl Deisseroth; Addgene plasmid #55639). We used EF1α promoter to drive the expression of target constructs for all AAV vectors except for AAV-DIO-mRuby2-T2A-Synaptophysin-eGFP, AAV-fDIO-hM4D(Gi)-mCherry, AAV-DIO-axon-GCaMP6s-P2A-mRuby3, and AAV-FLEX-oChIEF-mCitrine which are driven by the human synapsin1 promoter.

All AAV vectors used in this study were packaged as serotype DJ and generated as previously described^42^. In brief, AAV vectors were produced by transfection of AAV293 cells (Agilent) with three plasmids: an AAV vector plasmid carrying target constructs (DIO-mRuby2-T2A-Synaptophysin-eGFP, DIO-eGFP, fDIO-eGFP, fDIO-oPBG, fDIO-mRuby2-P2A-TVA, fDIO-hM4D(Gi)-mCherry, or FLEX-oChIEF-mCitrine), AAV helper plasmid (pHELPER; Agilent), and AAV rep-cap helper plasmid (pRC-DJ, gift from M. Kay). At 72 h post-transfection, the cells were collected and lysed by a repeated freeze-thaw procedure. Viral particles were then purified by an iodixanol step-gradient ultracentrifugation and subsequently concentrated using a 100-kDa molecular cutoff ultrafiltration device (Millipore). The genomic titer was determined by quantitative PCR. The AAV vectors were diluted in PBS to a working concentration of approximately 10^13^ viral particles/ml.

EIAV genomic vector plasmids were constructed from pEIAV-SIN6.1-CBGFPW (a gift from John Olsen; Addgene #44173) by replacing eGFP coding sequence with DNA fragments containing either DIO-FlpO or fDIO-Cre. RG-EIAV vectors were generated by a modified version of a published protocol^43^. Briefly, HEK293-T cells were transfected with three plasmids: an EIAV genomic vector (pEIAV-CAG-DIO-Flp or pEIAV-CAG-fDIO-Cre), a helper packaging plasmid (pEV53B; a gift from John Olsen), and a pseudotyping plasmid encoding fusion protein FuG-B2, an envelope protein carrying extracellular and transmembrane domain of rabies virus glycoprotein and conjugated to the cytoplasmic domain of vesicular stomatitis virus (a gift from Kazuto Kobayashi)^44^. At 72 h post-transfection, viral particles were harvested from the media by centrifugation using SureSpin630 swinging bucket rotor (Thermo Scientific) at 5,700 rpm and 16,200 rpm for 16 h and 2 h, respectively. EIAV viral particles were reconstituted from the pellets with PBS and immediately stored at −80°C.

Rabies virus was generated as previously described^45^ from a full-length cDNA plasmid containing all components of the virus except the coding sequence of the rabies glycoprotein which was replaced with eGFP (a gift from Karl-Klaus Conzelmann). In brief, B7GG cells were transfected with a total of five plasmids: four plasmids expressing the viral components pcDNA-SADB16N, pcDNA-SADB16P, pcDNA-SADB16L, pcDNA-SADB16G, and the rabies virus genomic vector. The virus-containing media was collected 3-4 days post-transfection and used for further amplification. Viral particles were harvested from the media by centrifugation using SureSpin630 rotor at 20,000 rpm for 2 h. Rabies viral particles were reconstituted from the pellets with PBS and immediately stored at −80°C. To generate EnvA-pseudotyped, glycoprotein-deleted rabies virus expressing eGFP (EnvA-RVΔG-eGFP), we used a modified version of a published protocol^45^. BHK-EnvA cells were grown in four 15-cm dishes to 50-60% confluency. The cells were transduced with the rabies virus expressing eGFP generated in the previous step. After 3 hours, the media was removed and the cells were rinsed with PBS, trypsinized for 5 min at 37°C, and isolated by pipetting repeatedly with a P1000 pipettor. Each dish was observed carefully under the microscope to ensure that the cells did not aggregate. The cells were then pelleted by centrifugation and plated in four new dishes. Trypsinization was repeated two more times at 9 and 21 hours after transduction to ensure complete removal of the original rabies glycoprotein. The virus-containing media was collected 4 days post-transduction and concentrated as described above. Plasmids expressing the rabies viral components, B7GG, BHK-EnvA, and HEK-TVA cells were gifts from Edward M. Callaway.

### Stereotaxic viral injections and optic fiber/cannula implantation

Mice were anesthetized with a mixture of ketamine (100 mg/kg) and dexmedetomidine (1 mg/kg) and placed on a stereotaxic frame (David Kopf Instruments). The body temperature was maintained with a heating pad during surgery and recovery from anesthesia. All stereotactic coordinates were derived from Paxinos and Franklin mouse brain atlas. Viruses were infused into the brain by using pulled glass micropipettes coupled with a syringe pump (PHD ULTRA; Harvard Apparatus) at a rate of 100 nl/min. For tracing experiment to visualize output regions of GPe PV neurons, 150 nl of AAV-FLEX-mRuby2-T2A-Synaptophysin-eGFP was injected into the GPe of *PV*^*Cre*^ mice (anteroposterior - 0.35 mm, mediolateral 1.95 mm from bregma, and depth −3.5 mm from brain surface). For cell-type- and projection-specific tracing, 500 nl of RG-EIAV-DIO-Flp was unilaterally injected into either SNr (anteroposterior −3.25 mm, mediolateral 1.6 mm from bregma, and depth −4.4 mm from brain surface) or PF (anteroposterior −2.15 mm, mediolateral 0.625 mm from bregma, and depth - 3.1 mm from brain surface) of *PV*^*Cre*^ mice along with 350 nl of AAV-fDIO-eGFP into ipsilateral GPe. After allowing 3 weeks of expression, mice were euthanized for circuit mapping analysis.

For mapping of input-output relationship, 500 nl of RG-EIAV-DIO-Flp was unilaterally injected into either SNr or PF of *PV*^*Cre*^ mice along with 350 nl of a 1:1 mixture of AAV-fDIO-oPBG and AAV-fDIO-mRuby2-P2A-TVA into ipsilateral GPe. After allowing 3 weeks of expression, animals were again anesthetized as previously described and 350 nl of RVΔG-eGFP(EnvA) was injected into ipsilateral GPe. Mice were euthanized after 7 days for quantification of input neurons and *in situ* hybridization.

For electrophysiological recordings of GPe neurons, 500 nl of RG-EIAV-fDIO-Cre was unilaterally injected into either SNr or PF of *PV*^*Flp*^ *x Ai14* mice. For recordings of postsynaptic neurons in SNr and PF, 350 nl of AAV-FLEX-oChIEF-mCitrine was unilaterally injected into the GPe of *PV*^*Cre*^ mice. After allowing 3 weeks of expression, acute slices were prepared for recordings.

For optogenetic behavioral experiments, *PV*^*Cre*^ mice were bilaterally injected with 350 nl of AAV-FLEX-oChIEF-mCitrine (or AAV-DIO-eGFP for controls) into the GPe. Subsequently, a bilateral 26-gauge guide cannula (cut length 4.5 mm, center-to-center distance 2 mm; Plastic One) was implanted over the medial forebrain bundle (MFB; anteroposterior −0.5 mm, mediolateral 1.0 mm from bregma) for 6-hydroxydopamine infusion and was temporally secured in place with cyanoacrylate glue. Optic fibers (200 μm, 0.22 NA; Doric Lenses) were cut to length (4.5 mm for SNr and 3.5 mm for PF) and implanted bilaterally above SNr (depth 4.0 mm from brain surface) or PF (mediolateral angle 15°, anteroposterior −2.15 mm, mediolateral −1.65 from bregma, depth −3.0 mm from brain surface) during the same surgery session. The optic fibers and the guide cannula were secured in place with a layer of adhesive cement (Radiopaque L-powder for C&B Metabond; Parkell). Once dried, absorbable sutures and a second layer of dental cement (Ortho-Jet; Lang Dental) were used to seal the head incision. For chemogenetic behavioral experiments, 500 nl of RG-EIAV-DIO-Flp was bilaterally injected into either SNr or PF of *PV*^*Cre*^ mice along with 350 nl AAV-fDIO-hM4D(Gi)-mCherry (or AAV-fDIO-eGFP for controls) into the GPe. During the same surgery session, a bilateral guide cannula was implanted over the MFB as described above. Behavioral experiments were performed 4 weeks after surgery to allow time for recovery and optimal viral expression.

For fiber photometry, *PV*^*Cre*^ mice were unilaterally injected with 350 nl of AAV-DIO-axon-GCaMP6s-P2A-mRuby3 into the GPe. Optic fibers (400 μm, 0.48 NA; Doric Lenses) were cut to length (4.5 mm for SNr and 3.5 mm for PF) and unilaterally implanted above ipsilateral SNr (depth 4.0 mm from brain surface) and PF in conjunction with a head bar during the same surgery session as described above. Behavioral experiments and recordings were performed 3 weeks after surgery to allow time for recovery and optimal viral expression.

Upon completion of behavioral experiments, viral injections and optic fiber placements were confirmed with histology to ensure proper targeting (Extended Data Fig. 3a-b and 5a-b).

### Dopamine depletion

6-hydoxydopamine (6-OHDA) solution was freshly prepared by adding 6-OHDA HCl (Sigma) to an appropriate volume of sterile saline (0.9% NaCl) with 0.02% sodium ascorbate (Sigma) to achieve the desired concentration. To prevent lesioning of non-dopamine monoaminergic neurons, desipramine (25 mg/kg) was injected intraperitoneally 30 min before 6-OHDA injection^46^.

6-OHDA injection for electrophysiological recordings was performed with the same method mentioned above two weeks after viral injection. Mice were anesthetized with isoflurane (2% in O_2_) and 750 nl of 6-OHDA solution (2.5 μg/μl) was unilaterally injected into the MFB (anteroposterior −0.5 mm, mediolateral 1.0 mm from bregma, depth −4.8 mm from brain surface) at a rate of 150 nl/min. For behavioral experiments after dopamine depletion, 2-3 weeks after viral injection and cannula implantation (see ‘behavioral assay’ for experimental timelines), mice were anesthetized with isoflurane (2% in O_2_) and a 33-gauge bilateral injector (cut length 5 mm; Plastic One) attached to a syringe pump (PHD ULTRA; Harvard Apparatus) was lowered into the guide cannula previously implanted over the MFB (depth −4.8 mm from brain surface). 750 nl of 6-OHDA solution (high-dose, 2.5 μg/μl for locomotion; low-dose, 1.25 μg/μl for reversal-learning task) was bilaterally infused into the MFB at a rate of 150 nl/min. Sham-operated control mice underwent similar procedures and were injected with vehicle (saline with 0.02% sodium ascorbate) instead of 6-OHDA. After infusion, the body mass of each animal was carefully monitored and a nutritionally fortified water gel (DietGel Recovery; ClearH_2_O) was provided in conjunction with a shallow water dish and moist mouse chow to aid recovery. Recordings and locomotion measurements were performed 7-10 days after 6-OHDA infusion. Reversal-learning task was performed 3 days after 6-OHDA infusion.

### SHIELD-MAP tissue processing

*PV*^*Cre*^ mice injected with AAV-FLEX-mRuby2-T2A-Synaptophysin-eGFP were processed for SHIELD-MAP as previously described^47^. In brief, mice were deeply anesthetized with isoflurane and were transcardially perfused with 10 ml of ice-cold PBS followed by 20 ml of freshly-made ice-cold SHIELD perfusion solution. The brains were carefully extracted and post-fixed in the same solution at 4°C for 48 hours. The brains were cut parasagittally into 3-mm blocks, containing the GPe, SNr, and PF and transferred to a new conical tube with 20 ml of ice-cold SHIELD Off solution and incubated at 4°C for 24 hours. Brain blocks were then incubated in SHIELD On solution at 37°C for 24 hours and washed in PBS containing 0.02% sodium azide (Sigma) overnight at room temperature (RT). Subsequently, the samples were cleared with SDS clearing buffer for 10 days at 45°C while shaking, then washed overnight at RT with PBS containing 1% (v/v) Triton-X 100 (Sigma) and then with PBS. The samples were then incubated in monomer solution at 4°C for 3 days and were gel-embedded under nitrogen gas purge at 33°C for 4 hours. The samples were rehydrated in PBS for several hours and transferred to deionized water for tissue expansion. A light-sheet microscope (SmartSPIM; Life Canvas Technologies) with a 10x, 0.6 NA objective (Olympus) was used to obtain images from expanded samples. Formulas for all solutions and a detailed protocol can be found at chunglabresources.com.

### Histology

After designated time for viral expression in tracing experiments or immediately after behavioral experiments, mice were deeply anesthetized with isoflurane and were transcardially perfused with ice-cold 4% paraformaldehyde in PBS. Brains were carefully extracted and post-fixed in the same fixative at 4°C for at least 12 hours. Brains were sliced at 60 μm thickness with a vibratome (VT1000; Leica). Brain sections were mounted on glass slides (SuperFrost Plus; Fisher Scientific) or collected and stored in cryoprotectant (30% ethylene glycol, 30% glycerol in PBS) at −20°C for immunohistochemistry. Mounted slides were coverslipped with DAPI Fluoromount-G (Southern Biotech) and imaged with a 10x objective on an Olympus VS120 virtual slide microscope.

### Tyrosine hydroxylase immunoreactivity

Quantification of striatal tyrosine hydroxylase (TH) immunoreactivity was used to assess the degree of dopamine depletion. Immunohistochemistry was carried out in free-floating fixed brain sections containing the dorsal striatum. The sections were washed with PBS for 3 times (10 min each) and incubated at room temperature for 1 hour in blocking solution containing 10% normal horse serum (Abcam), 0.2% bovine serum albumin (Sigma), and 0.5% Triton-X 100 (Sigma) in PBS. The sections were then incubated at 4°C for 20 hours in carrier solution containing 1% normal horse serum, 0.2% bovine serum albumin, and 0.5% Triton-X 100 in PBS with anti-TH primary antibody (1:2000; Millipore AB152). The sections were washed in PBS for 3 times and were incubated in carrier solution containing Alexa-flour 647-conjugated donkey anti-rabbit secondary antibody (1:1000; Life Technologies A-31573) at room temperature for 2 hours. Finally, the section washed 2 times in PBS, mounted, coverslipped, and imaged as described above. Each slide always included sections from naïve control animals that were processed and imaged in parallel to use as a reference. We used the pixel-intensity measuring tool in Fiji (ImageJ) to analyze fluorescence intensity from a 500 × 500 μm area taken from both hemispheres of each section. The fluorescence intensity for each section was normalized to the intensity of the naïve control tissue from the same slide. For locomotion measurements, all data included in the analysis were from mice with <35% TH immunoreactivity. For reversal-learning task, all data included in analysis were from mice with <70% TH immunoreactivity (Extended Data Fig. 7).

### Multiplex *in situ* hybridization

Mice that underwent surgeries for mapping of the input-output relationship were deeply anesthetized with isoflurane and transcardially perfused with ice-cold PBS. The brains were extracted, submerged in embedding medium (Tissue-Tek O.C.T.; Sakura), and frozen with 2-methylbutane (Sigma) chilled with dry ice in 70% ethanol. The frozen brain blocks were stored at −20°C for at least one day, then sliced with a cryostat microtome (Thermo-Fisher) to obtain 20-μm coronal sections. For each brain, 8 sections regularly sampled across the entire dorsal striatum were mounted on SuperFrost Plus slides (Fisher Scientific) and processed exactly as described in the RNAscope assay online protocol (ACD; Advanced Cell Diagnostics). We used probe against eGFP mRNA to label striatal neurons infected with EnvA-RVΔG-eGFP that spread trans-synaptically from either PV^GPe-SNr^ or PV^GPe-PF^ starter cells. Probes against *Drd1a* and *Drd2* mRNAs were used to determine whether the labeled striatal neurons were dMSNs or iMSNs. All probes were purchased from ACD. Because the labeling of eGFP mRNA was the strongest, we assigned the Atto-647 secondary probe to the eGFP probe to prevent bleed-through of the fluorescent signal into neighboring channels during imaging. Images of the entire dorsal striatum were acquired through a 30x silicone-immersion objective on a confocal microscope (Fluoview FV1200; Olympus). All eGFP-labeled neurons were manually quantified and classified as either dMSN or iMSN.

### Quantification of input-output relationship

All sections except the olfactory bulb and the cerebellum were collected and processed as described in the ‘Histology’ section. All eGFP-labeled neurons from all brain regions except the GPe (injection site of the rabies virus) were quantified and presented as a percentage of total eGFP-labeled neurons from each brain. Brain regions were assigned based on the Franklin and Paxinos Mouse Brain Atlas. Regions with <1% of total eGFP-labeled neurons were not included in the summary plot in Fig. 1.

### *Ex vivo* electrophysiology

Mice were anesthetized with isoflurane and transcardially perfused with ice-cold choline-based slicing solution, containing (in mM): 25 NaHCO_3_, 1.25 NaH_2_PO_4_, 2.5 KCl, 7 MgCl_2_, 25 glucose, 0.5 CaCl_2_, 110 choline chloride, 11.6 sodium ascorbate, and 3.1 sodium pyruvate. Brains were carefully extracted and transferred to a chamber filled with the same solution on a vibratome (VT1200; Leica). Brains were sliced at 250 μm (coronally for recording of SNr and PF neurons, parasagittally for PV^GPe-SNr^ neurons, and horizontally for PV^GPe-PF^ neurons) and incubated at 35°C for 15-20 min in recovery solution, containing (in mM): 118 NaCl, 2.6 NaHCO_3_, 11 glucose, 15 HEPES, 2.5 KCl, 1.25 NaH_2_PO_4_, 2 sodium pyruvate, 0.4 sodium ascorbate, 2 CaCl_2_, and 1 MgCl_2_. Slices were maintained at room temperature for at least one hour until transferred to a recording chamber an Olympus BX51WI upright microscope. The chamber was continuously superfused with artificial cerebrospinal fluid (ACSF), containing (in mM): 125 NaCl, 25 NaHCO_3_, 2.5 KCl, 1.25 NaH_2_PO_4_, 11 glucose, 1.3 MgCl_2_, and 2.5 CaCl_2_, maintained at 30 ± 2°C by a feedback temperature controller. Slicing solution, recovery solution, and ACSF were constantly bubbled with 95% O_2_ and 5% CO_2_. All compounds were purchased from Tocris or Sigma.

For all recordings, patch pipettes (3-5 MΩ) were pulled from borosilicate glass (G150TF-4; Warner Instruments) with a DMZ Universal Electrode Puller (Zeitz Instruments) and filled with appropriate intracellular solutions. Liquid junction potential was not corrected for any experiments. Neurons were visualized with differential interference contrast optics or epifluorescence (Olympus). Recordings were made with a MultiClamp700B amplifier and pClamp10 software (Molecular Devices). Data were low-pass filtered at 1 kHz and digitized at 10 kHz with a digitizer (Digidata 1440; Molecular Devices). Series resistance was monitored and cells that displayed > 20% change over the duration of recording were excluded.

For current-clamp recording, pipettes were filled with an intracellular solution containing (in mM): 125 K^+^-gluconate, 4 NaCl, 10 HEPES, 0.5 EGTA, 20 KCl, 4 Mg^2+^-ATP, 0.3 Na^+^-GTP, and 10 Na_2_-phosphocreatine (290-300 mOsm, pH 7.2). Autonomous firing activity was acquired in a cell-attached configuration for 1 min prior to break-in. To measure the firing capacity of identified PV^GPe-SNr^ and PV^GPe-PF^ neurons, baseline firing was maintained at 5Hz and currents of increasing intensity (20 pA increments, duration = 500 ms) were injected until neurons went into depolarization block. All current-clamp recordings were performed in the presence of 5 μM NBQX and 50 μM picrotoxin to block synaptic transmission. Detection of spikes and measurements of action potential (AP) characteristics were performed using a custom Python script with the following criteria. AP threshold was measured as a change in voltage from rest at which the slope = 20 mV/ms. Peak amplitude was measured as a change in voltage from AP threshold to the peak of AP, and afterhyperpolarization (AHP) as a change in voltage and time from threshold to minimum after the peak. Half-width was calculated as full-width at half max amplitude. All measurements were quantified while neurons were firing at 5Hz.

Passive membranes properties were calculated from voltage-clamp recordings with a custom Python script based on previously described method^48^; membrane resistance (*R*_m_) was calculated by *R*_m_ = Δ*V*_test_/Δ*I* where Δ*V*_test_ is a 10-mV step (50 ms), and Δ*I* is the difference between steady-state current and baseline during the last 10 ms of the voltage step. Membrane capacitance (*C*_m_) was calculated by *C*_m_ = *Q*_t_ * Δ*V*_test_ where *Q*_t_ is the integral of the transient current elicited by Δ*V*_test_, a 10-mV voltage step (50 ms).

To measure the ratio of excitatory and inhibitory inputs (E/I ratio), pipettes were filled with a Cs-based intracellular solution, containing (in mM): 115 Cs^+^-methanesulphonate, 10 HEPES, 1 EGTA, 1.5 MgCl2, 4 Mg^2+^-ATP, 0.3 Na^+^-GTP, 10 Na_2_-phosphocreatine, 2 QX 314-Cl, 10 BAPTA-tetracesium (295 mOsm, pH 7.35). Electrically-evoked excitatory postsynaptic and inhibitory postsynaptic currents (EPSCs and IPSCs, respectively) were recorded from identified PV^GPe-SNr^ and PV^GPe-PF^ neurons at −60 mV (for EPSCs) and 0 mV (for IPSCs). E/I ratio was calculated by dividing the amplitude of EPSC by the amplitude of IPSC. Spontaneous EPSC and IPSC were also recorded at −60 mV and 0 mV, respectively.

For the recording of Sr^2+^-induced asynchronous optogenetically-evoked postsynaptic currents, pipettes were filled with high-chloride intracellular solution, containing (in mM): 122 CsCl, 8 NaCl, 10 glucose, 1 CaCl_2_, 10 HEPES, 10 EGTA, 2 Mg^2+^-ATP, 0.3 Na^+^-GTP, 2 QX 314-Cl (280-290 mOsm, pH 7.2). Slices were incubated with Ca^2+^-free ACSF containing 4 mM SrCl_2_ for 30 min before recording. oChIEF-expressing axon terminals were stimulated with a 5-ms blue light pulse emitted from a collimated light-emitting diode (473 nm; Thorlabs) driven by a T-Cube LED Driver (Thorlabs) under the control of Digidata 1440A Data Acquisition System and pClamp10 software (Molecular Devices). Light was delivered through the reflected light fluorescence illuminator port and the 40X objective at maximum intensity (13.45 mW). Recordings of optogenetically-evoked IPSCs were obtained from SNr or PF neurons located close to mCitrine-labeled axons in the presence of 5 μM NBQX. IPSC events were analyzed using a custom Python script based on a previously described method^49^; the best fit curve of the largest evoked transient was subtracted from the original trace and the amplitude and frequency of IPSC events were measured between 0-400 ms after photostimulation.

To measure paired-pulse ratio (PPR), pipettes were filled with a Cs-based intracellular solution. Optogenetically-evoked IPSCs were recorded from SNr or PF neurons located close to mCitrine-labeled axons while the GPe-PV axons were stimulated with two 5-ms light pulses separated by a 100-ms interval. PPR was calculated as the ratio of 2^nd^ IPSC peak amplitude / 1^st^ IPSC peak amplitude. Target recordings in the SNr (Extended Data Fig. 5c-d) was performed similarly with trains of 5, 10, 20, 50 Hz light pulses, or a constant 500-ms light pulse.

### Behavioral assay: locomotion

Locomotion measurements in freely moving mice were performed in an open field arena (opaque white acrylic; w: 30 cm, L: 30 cm, H: 30 cm). To acclimate mice to the arena, we allowed them to freely explore the arena for 1 hour per day for 3 consecutive days. Baseline locomotor activity was obtained on the next day from 10-60 min after start during a 1-hour session. On subsequent days, the mice were tested based on their behavioral manipulation. The activity was recorded as a movie from overhead with a 15 Hz frame rate. The center point of each mouse in each frame was tracked off-line using Viewer II software (Biobserve) and was exported as raw data. Locomotor activity was then analyzed using a custom Python script to calculate speed in centimeter per second.

### Behavioral assay: reversal-learning task

Mice were tested on a modified version of naturalistic foraging reversal discrimination task originally developed for rats^50,51^ and adapted for mice^52,53^. Mice were food deprived to reduce body weight to 80-85% of the *ad libitum* feeding weight and habituate with a small animal food bowl (Lixit Nibble Food Bowl) used during the task for 7 days in their home cage prior to testing. During the same 7-day period, each mouse was handled for 1 min per day to minimize stress associated with handling during testing. One day before testing, mice were acclimated to a testing chamber (opaque white acrylic; w: 30 cm, L: 60 cm, H: 30 cm). Each mouse was placed in a waiting area separated from the rest of the testing chamber by a removable start gate. At the start of each trial, the gate was lifted and the mouse was allowed to explore two bowls placed at the opposite end of the chamber until a food reward was found and consumed. Both bowls did not contain digging media and only one bowl contains a food reward (a piece of Honey Nut Cheerios ∼50 mg).

The task consists of two consecutive phases that were performed over a 2-day period. First, in the association phase, mice were trained to associate a certain cue with a rewarding outcome, and a different cue without reward until they can discriminate reliably. In the subsequent reversal-learning phase, the cue-outcome associations were reversed, and thus, the mice needed to switch their actions to obtain the reward. The cues were either olfactory (odor) or somatosensory (texture of the digging medium which hides the bait). When the olfactory cues were used, one bowl contained coconut-scented soft bedding (National Geographic) and the other contained lavender-scented soft bedding (National Geographic). When the somatosensory cues were used, one bowl contained aquarium sand (National Geographic) and the other contained aquarium gravel. Only one type of cue was used with each mouse for the entire period of the task. All cues were presented in identical small animal food bowls that are identical in color and size. Digging media were mixed with the Honey Nut Cheerios powder (0.1% by volume).

On the first testing day (association), during each trial, the mouse was kept in the waiting area and two bowls were present at the opposite end of the chamber separated from the mouse by the start gate. At the start of each trial, the gate was lifted and the mouse is allowed to explore two bowls until digging in one bowl to signify a choice. The food reward was paired with a certain cue throughout the task and was hidden in the digging medium. The baited bowl was pseudorandomly presented on either side of the testing cage. The mouse was required to choose the correct cue that would result in getting a food reward. The association phase was complete when a criterion of eight correct trials in a block of ten trials is met (80% accuracy). On the second day (reversal learning), the mouse was presented with the same set of cues as in the association phase but the cue-reward pairing is reversed. Similar criterion was required to complete the reversal-learning phase. For analysis of event-locked activity in Fig. 3c-f, reversal-learning phase was broken down into: early reversal learning (all trials before reaching the last ten-trial block, representing a period before the criterion was met), and late reversal learning (all trials in the last ten-trial block, representing a period at which the criterion was met).

The number of correct and incorrect choice during each phase was scored manually and the performance of each mouse was reflected by the number of trials needed per phase to meet the criterion. Similar to previous findings, we observed an increase in the number of trials to criterion if odor is the relevant cue^54^. Therefore, performance data for each mouse was normalized to an average number of trials to criterion from all naïve control mice (control mice that did not receive photostimulation or CNO) that were presented with a similar type of cue (odor or somatosensory) during the task. In addition, error analysis was performed in the reversal-learning phase to measure the ability to maintain new cue-reward association after switching. Incorrect trials were classified as perseverative errors when the incorrect bowl was chosen more than 3 times in a block of 4 consecutive trials. Once a mouse started making fewer than 3 errors in a block, all subsequent incorrect trials were considered regressive errors^55^.

### Fiber photometry

GCaMP6s and mRuby3 fluorescence was collected simultaneously at 20Hz through a patchcord of bundled 400-μm diameter, 0.48 NA fibers (Doric Lenses) coupled to a setup similar to that previously described^56^. We used Lumencor SPECTRA X Light Engine as an excitation light source for cyan (470 nm) and green (550 nm) light. At the start of each recording session, light intensity was adjusted to 120 μW (measured at the tips of the patchcord). Photometry data were analyzed with a custom Python script. First, data from the red channel were scaled to the green channel using a linear fit and were used as baseline (F_0_) to correct for motion artifacts. ΔF was then calculated by subtracting this baseline. ΔF/F_0_ was then smoothed by filtering with a 2^nd^ order Savitzky-Golay filter. Finally, ΔF/F_0_ from each recording session was z-scored relative to the entire session and aligned to behavioral events.

To determine the relationship of fluorescence signals to locomotion, recordings were performed while the mice were head-fixed on a cylindrical treadmill (15-cm diameter). The treadmill velocity was sampled at 20Hz by a rotary encoder (TRD-SH Series; AutomationDirect) coupled to the axel of the treadmill. An Arduino was used to send triggers to the sCMOS camera (Orca-Flash4; Hamamatsu) to synchronize the treadmill velocity and fluorescence data stream. Locomotion initiation was identified as the time point at which the treadmill acceleration was greater than 10 cm/s^2^ and was not preceded within by 1-s window with an average velocity lower than 1 cm/s, indicating a clear transition between rest and treadmill locomotion. Termination was identified as the time point at which the treadmill acceleration was lower than −10 cm/s^2^ and the average velocity in the 1-s window succeeding that time point was lower than 1 cm/s, indicating a rapid transition from locomotion to rest. Z-scored ΔF/F_0_ from each recording session was aligned to locomotion initiations and terminations. These traces were then averaged across all sessions and mice to generate summary traces.

In the reversal-learning task, recordings were performed while the mice were preforming the reversal-learning task as described above. The following behavioral time points were manually marked off-line from the videos recorded during the task: trial start, choice (the moment the mouse started digging), and trial end. Z-scored ΔF/F_0_ from each recording session was aligned to these behavioral time points. These traces were then averaged across all mice to generate summary traces. An average of z-scored ΔF/F_0_ was calculated from 0-2 s after behavioral onsets to compare between different stages of the task.

### Optogenetic stimulation

A 473 nm blue laser diode (OEM Laser Systems) was connected to the ferrule of the optic fiber previously implanted on mice though a plastic sleeve. Laser power was measured before each experiment and adjusted to ∼8 mW. For all experiments with optogenetic activation, photostimulation consisted of 500-ms pulses repeated every 1.5 s. The stimulation paradigm was determined by testing the efficacy of a range of stimulation frequencies that can reliably inhibit spiking activity of SNr neurons *ex vivo* and was used throughout this study (Extended Data Fig. 5c-d). Light pulses were generated with a TTL pulse generator (OPT_G4; Doric Lenses). For locomotion assay, light pulses were delivered at for 10 min starting at 30 min after session start during a 1-hour session (Fig. 4g-h). For the reversal-learning task, light pulses were delivered for the entire duration of each trial during the reversal-learning phase.

### Chemogenetic inhibition

Clozapine-*N*-oxide (CNO; Enzo) was dissolved in water to obtain a stock concentration of 5 mg/ml and stored in small aliquots at −20°C. The working solution was freshly prepared before each use by diluting the stock with 0.9% saline to obtain a concentration of 0.5 mg/ml. Mice were injected with 0.01 ml/g body weight of the working solution to achieve a dose of 5 mg/kg. For locomotion assay, on the first test day, mice were injected with saline 1 hour before testing. On the second day, mice were injected with the working solution 1 hour before testing. For the reversal-learning task, mice were injected with the working solution 1 hour before the first trial of the reversal-learning phase.

### Statistics

Statistical analyses were performed using Prism 8 (Graphpad Software). All statistical data can be found in figure legends with corresponding sample sizes. The sample sizes were chosen based on common practice in mouse behavioral experiments. All data were tested for normality with Shapiro-Wilk test. Then the appropriate parametric or non-parametric tests were applied. Correction for multiple comparisons was performed using the Bonferroni method. All statistical tests were two-tailed. Statistical significance levels were set at \**p* < 0.05, \*\**p* < 0.01, \*\*\**p* < 0.001. All data are presented as mean ± SEM.

## Code availability

The code that supports the findings of this study is available from the corresponding author upon request.

## Data availability

The data that support the findings of this study are available from the corresponding author upon request.

