## Supplementary material for "Divergent pallidal pathways underlying distinct Parkinsonian behavioral deficits": Supple

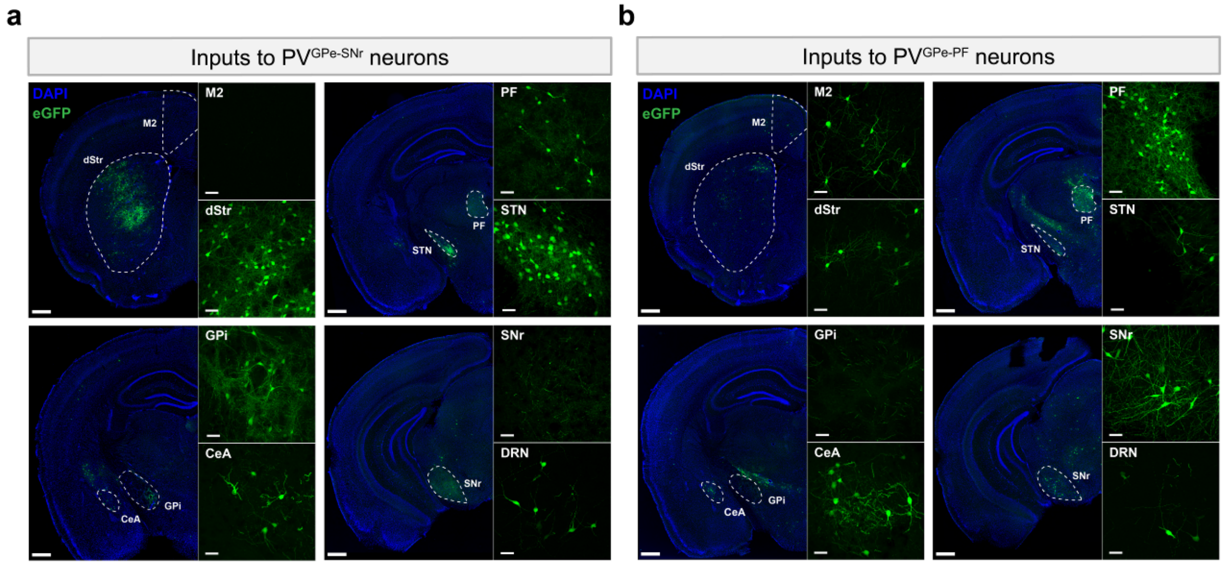


**Extended Data Fig. 1 | Whole-brain mapping of inputs to PV$^{GPe-SNr}$ and PV$^{GPe-PF}$ neurons.** Representative images of select brain areas showing transsynaptically labeled input neurons of PV$^{GPe-SNr}$ neurons. **b**, same as in **a**, but for PV$^{GPe-PF}$ neurons. M2, secondary motor cortex; dStr, dorsal striatum; CeA, central amygdala; DRN, dorsal raphe nucleus. Scale bars, 500 µm; inset 50 µm.

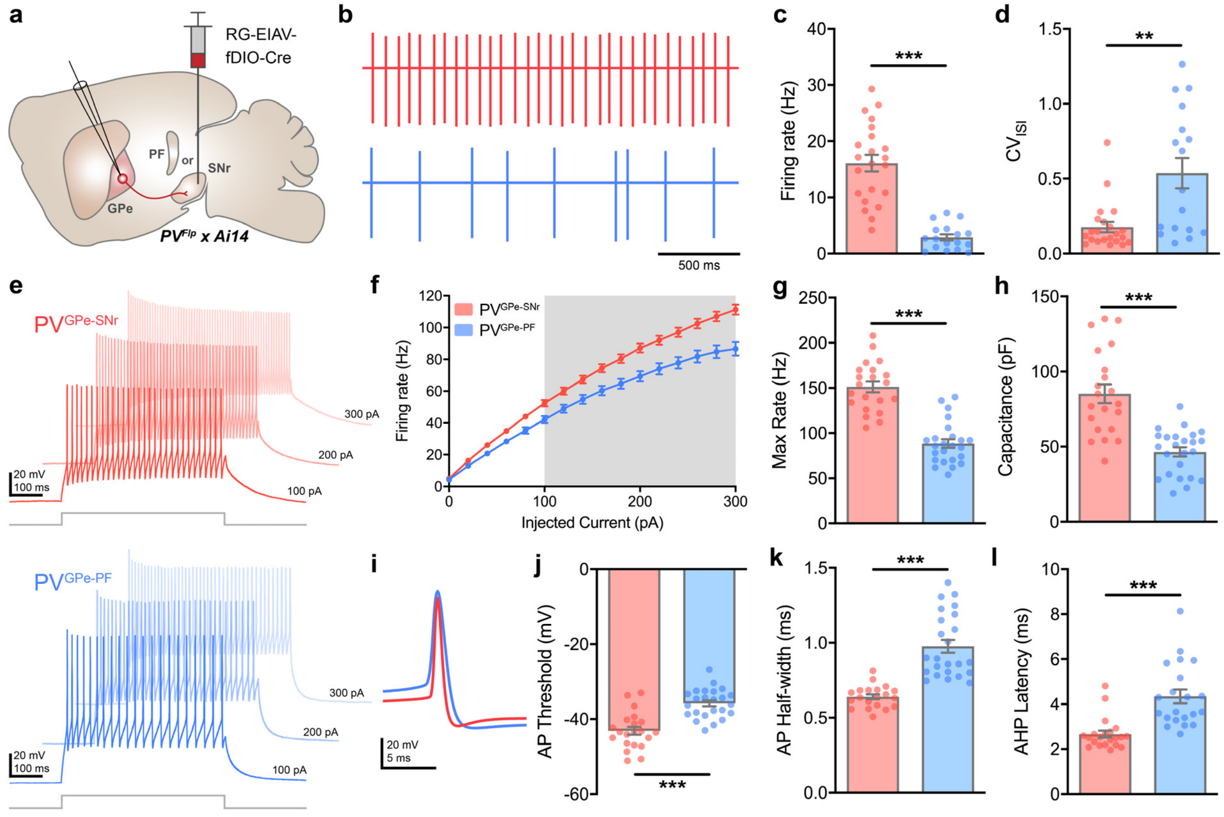


**Extended Data Fig. 2 | PV$^{GPe-SNr}$ and PV$^{GPe-PF}$ neurons exhibit distinct electrophysiological properties.** **a**, Strategy using *PV$^{Flp}$ x Ai14* mouse for labeling GPe PV neurons in projection-specific manner. RG-EIAV-fDIO-Cre is injected into either SNr or PF. **b**, Representative cell-attached recordings from PV$^{GPe-SNr}$ and PV$^{GPe-PF}$ neurons. **c**, Autonomous firing rate of PV$^{GPe-SNr}$ and PV$^{GPe-PF}$ neurons. Mann-Whitney *U*-test, U = 7; ***$p < 0.001$ ($n = 22$ cells from 8 mice for PV$^{GPe-SNr}$ and $n = 17$ cells from 7 mice for PV$^{GPe-PF}$ neurons). **d**, Firing regularity of PV$^{GPe-SNr}$ and PV$^{GPe-PF}$ neurons. Mann-Whitney *U*-test, U = 7; **$p = 0.0019$ ($n = 22$ cells for PV$^{GPe-SNr}$ and $n = 17$ cells for PV$^{GPe-PF}$ neurons). **e**, Representative traces of neuronal firing in response to 100pA, 200pA, and 300pA current injections in PV$^{GPe-SNr}$ and PV$^{GPe-PF}$ neurons. **f**, Action potential firing frequency in response to a range of current injections for PV$^{GPe-SNr}$ and PV$^{GPe-PF}$ neurons. Gray shading shows a significant difference ($p < 0.05$). Multiple *t*-test, corrected for multiple comparisons using Holm-Sidak method ($n = 21$ cells from 7 mice for PV$^{GPe-SNr}$ and $n = 24$ cells from 11 mice for PV$^{GPe-PF}$ neurons). **g**, Maximum firing rate of PV$^{GPe-SNr}$ and PV$^{GPe-PF}$ neurons. Mann-Whitney *U*-test, U = 23; ***$p < 0.001$ ($n = 21$ cells for PV$^{GPe-SNr}$ and $n = 24$ cells for PV$^{GPe-PF}$ neurons). **h**, Membrane capacitance of PV$^{GPe-SNr}$ and PV$^{GPe-PF}$ neurons. Mann-Whitney *U*-test, U = 23; ***$p < 0.001$ ($n = 21$ cells for PV$^{GPe-SNr}$ and $n = 24$ cells for PV$^{GPe-PF}$ neurons). **i**, Representative traces of action potential waveforms recorded from PV$^{GPe-SNr}$ and PV$^{GPe-PF}$ neurons. **j**, Action potential threshold of PV$^{GPe-SNr}$ and PV$^{GPe-PF}$ neurons. Mann-Whitney *U*-test, U = 57; ***$p < 0.001$ ($n = 21$ cells for PV$^{GPe-SNr}$ and $n = 24$ cells for PV$^{GPe-PF}$ neurons). **k**, Action potential half-width of PV$^{GPe-SNr}$ and PV$^{GPe-PF}$ neurons. Mann-Whitney *U*-test, U = 10; ***$p < 0.001$ ($n = 21$ cells for PV$^{GPe-SNr}$ and $n = 24$ cells for PV$^{GPe-PF}$ neurons). **l**, Afterhyperpolarization latency of PV$^{GPe-SNr}$ and PV$^{GPe-PF}$ neurons. Mann-Whitney *U*-test, U = 32; ***$p < 0.001$ ($n = 21$ cells for PV$^{GPe-SNr}$ and $n = 24$ cells for PV$^{GPe-PF}$ neurons). All data presented as mean ± SEM.

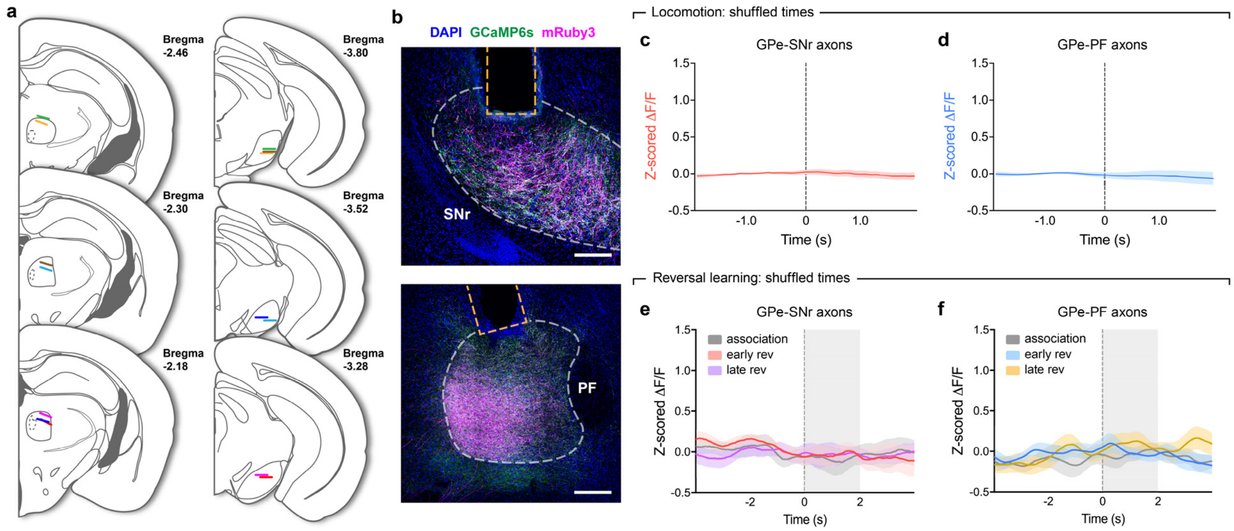


**Extended Data Fig. 3 | Controls for photometry recordings. a**, Optic fiber placements in PF and SNr for fiber photometry recordings. Different colors denote fibers from the same mouse (*n* = 7 mice). **b**, Representative images showing mRuby3 and axon-GCaMP6s expression in axons of GPe-PV neurons at implantation sites for optic fibers. Scale bars, 200 µm. **c**, Z-scored ΔF/F (averaged across all events) representing the activity of $PV^{GPe\text{-}SNr}$ axons at randomly chosen time points during treadmill locomotion. Number of events was determined based on the number of locomotion onsets in each recording session (*n* = 135 events). **d**, same as in **c**, but showing the activity of $PV^{GPe\text{-}PF}$ axons. **e**, Z-scored ΔF/F (averaged across 7 mice) representing the activity of $PV^{GPe\text{-}SNr}$ axons at randomly chosen time points during different stages of reversal-learning task. Number of events was determined based on the number of trials in each stage of the task. **f**, same as in **e**, but showing the activity of $PV^{GPe\text{-}PF}$ axons. Shaded areas accompanying the z-scored ΔF/F traces in **c-f** indicate SEM.

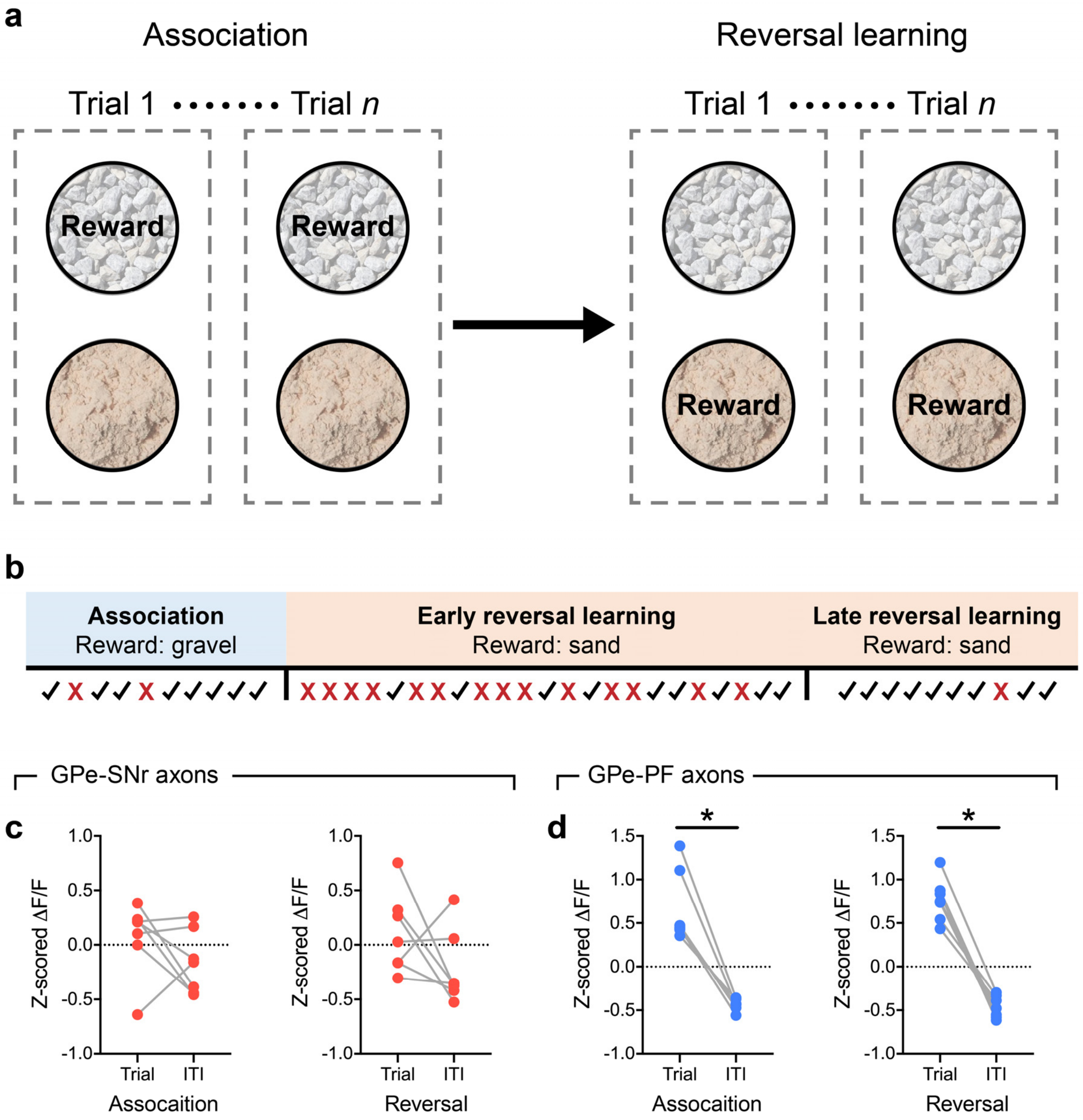


**Extended Data Fig. 4 | Structure of the reversal-learning task and comparison of overall neural activity during trials and inter-trial intervals.** **a**, Example task structure using gravel and sand as digging media to provided two different contexts. The reward is paired with sand in the association phase and is later switched to gravel in the reversal-learning phase. **b**, Timeline of the task showing how the association, early, and late reversal-learning stages are defined for a representative mouse. Check marks and crosses represent correct and incorrect trials, respectively. **c**, Comparison of mean z-scored ΔF/F for $PV^{GPe\text{-}SNr}$ axons between trial periods and inter-trial intervals (ITI) during the association phase (left; Wilcoxon sign-rank test, W = -12; $p = 0.3750$, $n = 7$ mice) and reversal-learning phase (right; Wilcoxon sign rank test, W = -18; $p = 0.1562$, $n = 7$ mice). **d**, same as in **c**, but for $PV^{GPe\text{-}PF}$ axons during the association phase (left; Wilcoxon sign-rank test, W = -28; $*p = 0.0156$, $n = 7$ mice) and reversal-learning phase (right; Wilcoxon sign rank test, W = -28; $*p = 0.0156$, $n = 7$ mice).

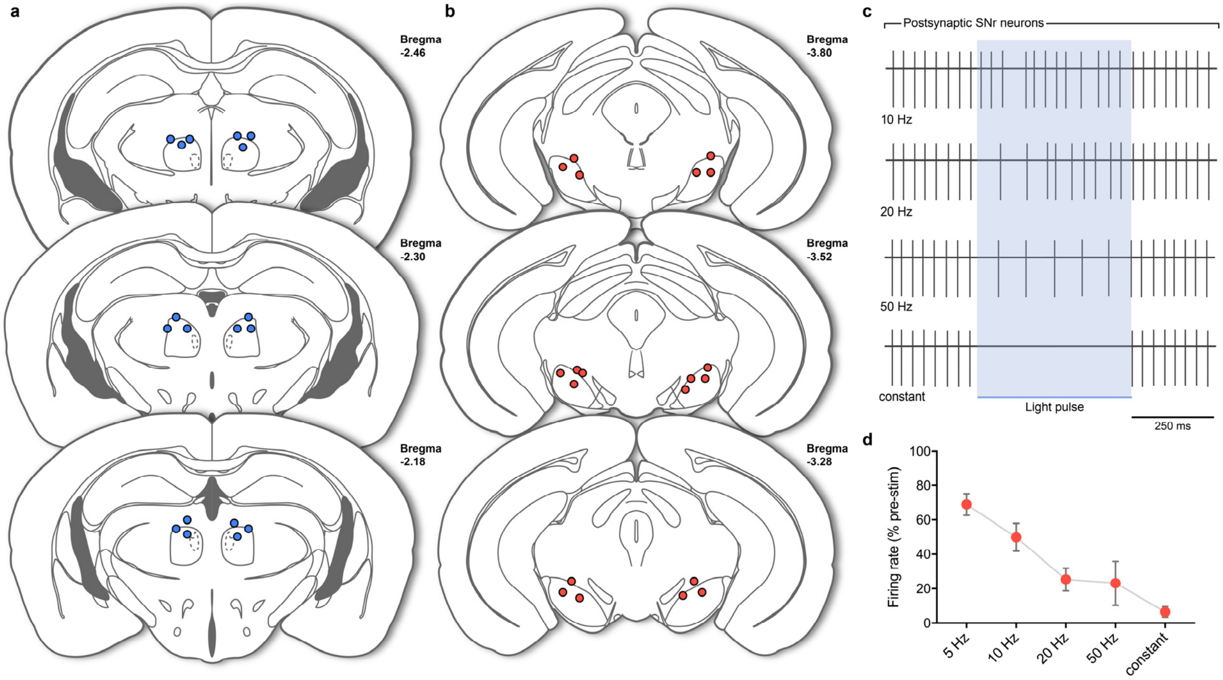


**Extended Data Fig. 5 | Activation of GPe-SNr synapses fully inhibits SNr neurons only when stimulated at high frequency. a**, Fiber tip locations in PF for data in Fig. 3i, 3l, 4j. and Extended Data Fig. 4c-d. **b**, Fiber tip locations in SNr for data in Fig. 3h, 3k, 4i, 5e, and Extended Data Fig. 4a-b, 7b, 7d-e. **c**, Representative traces from cell-attached recording showing inhibition of firing activity in SNr neurons during photostimulation of $PV^{GPe\text{-}SNr}$ axons at different frequencies. **d**, Firing rates of SNr neurons during 5-50 Hz and constant photostimulation of $PV^{GPe\text{-}SNr}$ axons ($n$ = 21 cells). Data presented as mean ± SEM.

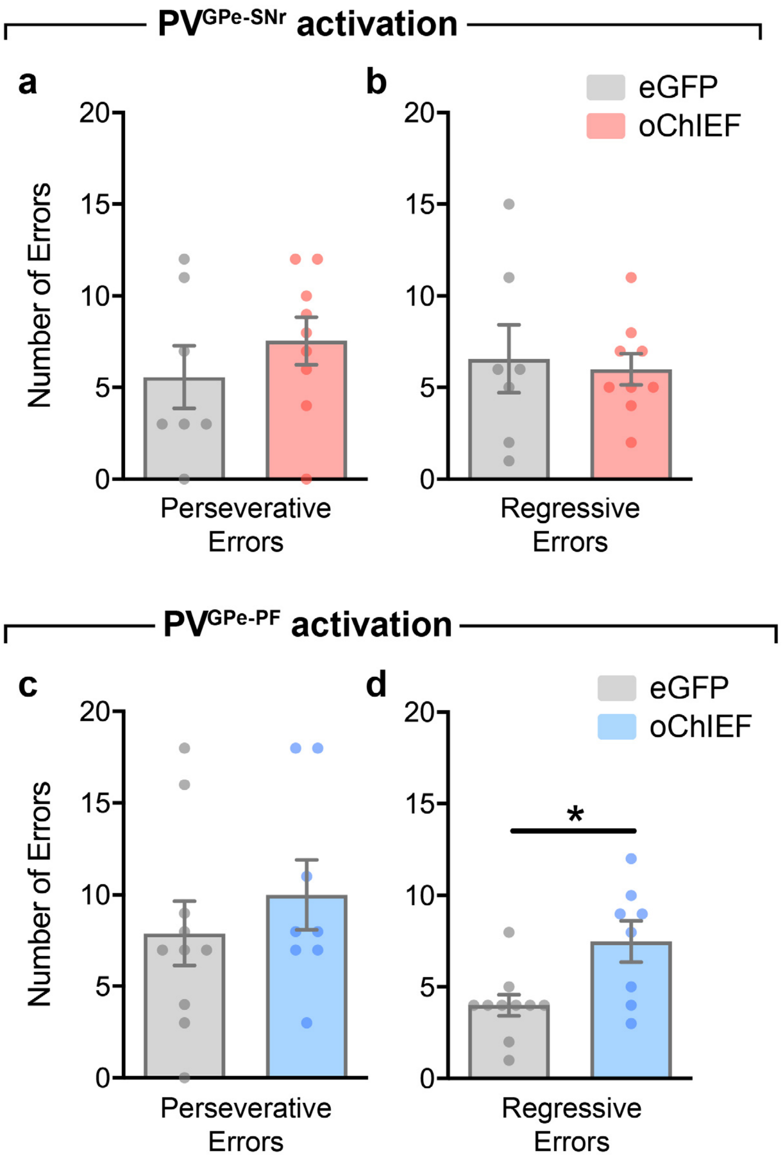


**Extended Data Fig. 6 | Activation of PV$^{GPe\text{-}PF}$ neurons increased number of regressive errors made during reversal learning. a-b**, Number of errors during the reversal-learning phase made by mice that received photostimulation in PV$^{GPe\text{-}SNr}$ neurons ($n$ = 7 mice for eGFP, $n$ = 9 mice for oChIEF)**. a**, Perseverative errors; Unpaired $t$-test, $t(14) = 0.9432$, $p = 0.3616$. **b**, Regressive errors; Unpaired $t$-test, $t(14) = 0.3002$, $p = 0.7684$. **c-d**, Number of errors during the reversal-learning phase made by mice that received photostimulation in PV$^{GPe\text{-}PF}$ neurons ($n$ = 10 mice for eGFP, $n$ = 8 mice for oChIEF)**. c**, Perseverative errors; Unpaired $t$-test, $t(16) = 0.8109$, $p = 0.4293$. **d**, Regressive errors; Unpaired $t$-test, $t(16) = 2.951$, $**p = 0.0094$. All data presented as mean ± SEM.

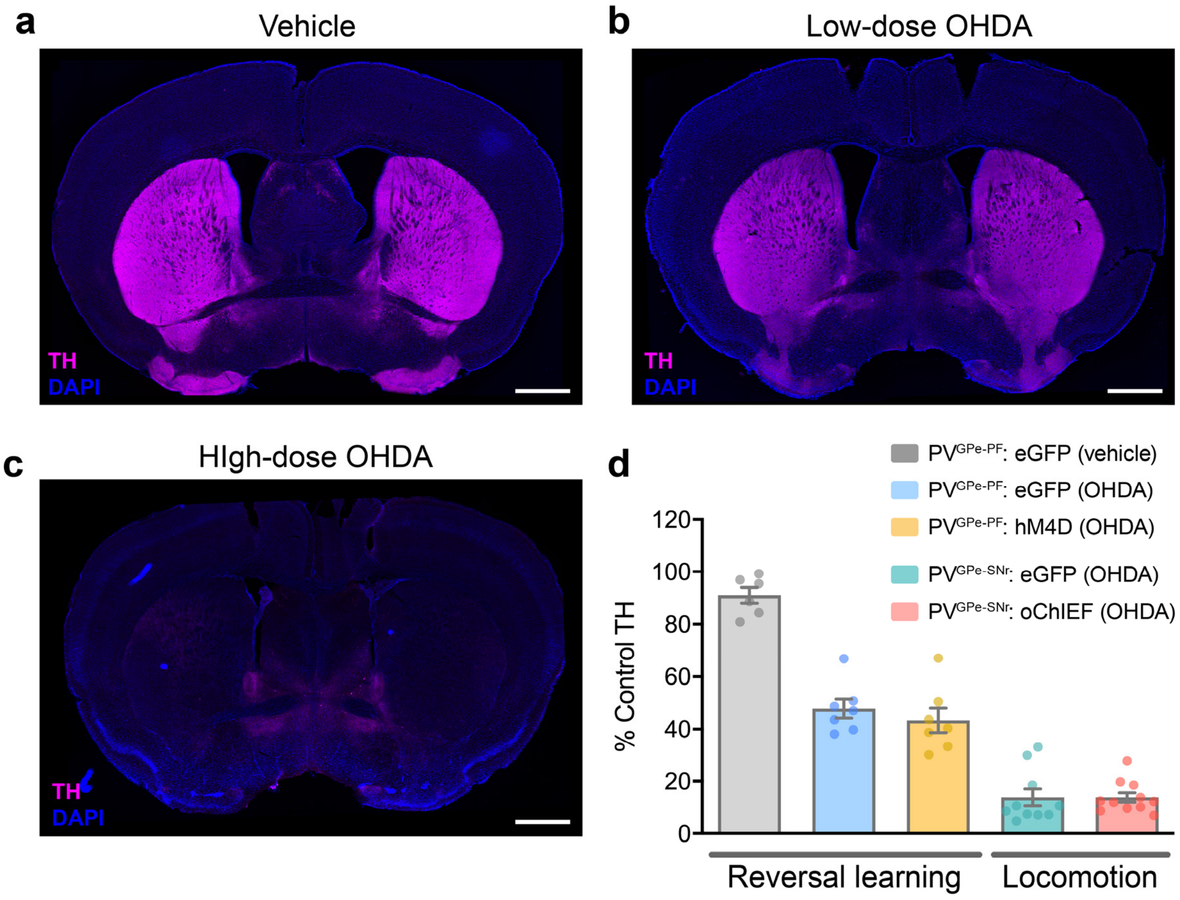


**Extended Data Fig. 7 | Quantification of TH immunoreactivity. a**, Representative image of TH immunoreactivity in the striatum of a mouse injected with vehicle (0.02% sodium ascorbate in 0.9% saline). **b**, Representative image of TH immunoreactivity in the striatum 3 days after bilateral injection of low-dose 6-OHDA (1.25 µg/µl). **c**, Representative image of TH immunoreactivity in the striatum 10 days after bilateral injection of high-dose 6-OHDA (2.5 µg/µl). **d**, Quantification of TH immunoreactivity at different stages of dopamine depletion in rescue experiments for reversal learning and locomotion. Data presented as % mean ± SEM of naïve control striatal sections. Scale bar, 1 mm.

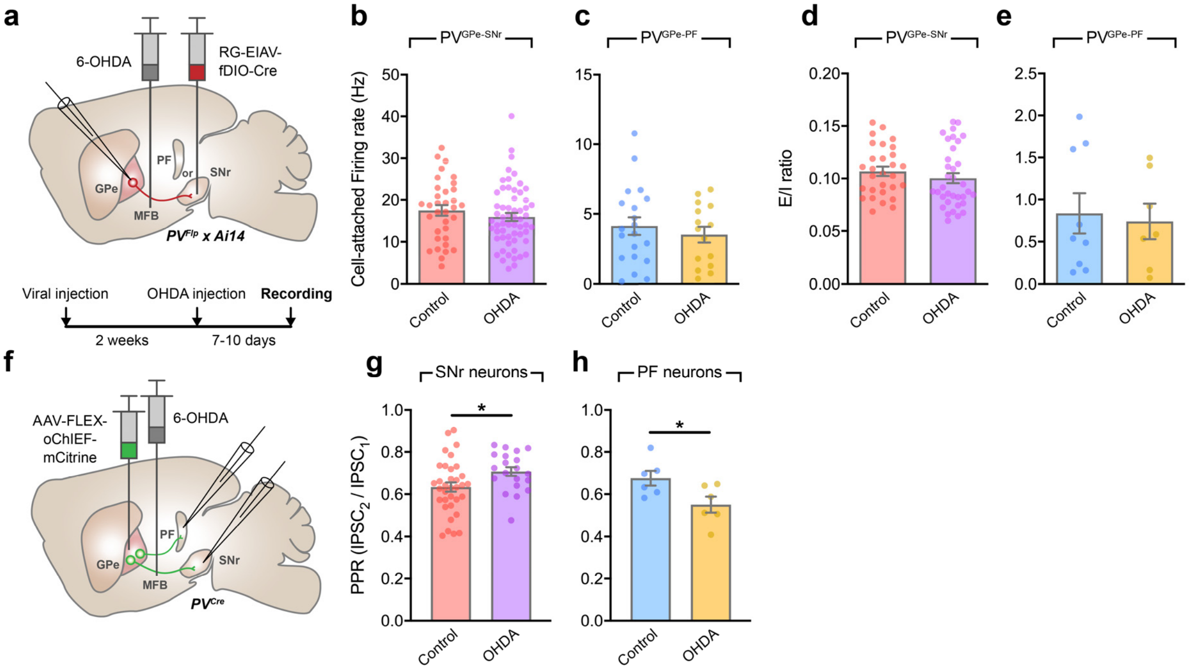


**Extended Data Fig. 8 | PV$^{GPe\text{-}SNr}$ and PV$^{GPe\text{-}PF}$ neurons exhibit distinct electrophysiological adaptations to dopamine depletion. a**, Schematic of viral and 6-OHDA injections and the experimental timeline for the recording of PV$^{GPe\text{-}SNr}$ and PV$^{GPe\text{-}PF}$ neurons in acute slices after dopamine depletion. Autonomous firing rate in PV$^{GPe\text{-}SNr}$ neurons (**b**; control $n$ = 34 cells from 9 mice, OHDA $n$ = 59 cells from 12 mice) and PV$^{GPe\text{-}PF}$ neurons (**c**; control $n$ = 20 cells from 8 mice, OHDA $n$ = 15 cells from 7 mice) recorded in cell-attached configuration in the presence of synaptic transmission blockers, NBQX and picrotoxin (PTX) in extracellular solution. Mann-Whitney *U*-test, U = 849; $p$ = 0.2213 in **b**, and U = 139; $p$ = 0.7297 in **c**. **d**, Ratio of evoked excitatory and inhibitory inputs onto PV$^{GPe\text{-}SNr}$ neurons (**d**; control $n$ = 29, OHDA $n$ = 35 cells) and PV$^{GPe\text{-}PF}$ neurons (**e**; control $n$ = 7, OHDA $n$ = 9 cells) from naïve control and dopamine-depleted mice. Mann-Whitney *U*-test, U = 419; $p$ = 0.2371 in **d**, and U = 29; $p$ = 0.8371 in **e**. **f**, Schematic of viral and 6-OHDA injections for the recording of SNr and PF neurons in acute slices after dopamine depletion. The release probability at GPe-SNr and GPe-PF synapses were altered in dopamine-depleted mice (**g-h**). Paired-pulse ratio (PPR; 2$^{nd}$ IPSC peak / 1$^{st}$ IPSC peak amplitude) measured from SNr neurons (**g**; control $n$ = 34 cells from 8 mice, OHDA $n$ = 20 cells from 6 mice) and PF neurons (**h**; control $n$ = 6 cells from 3 mice, OHDA $n$ = 6 cells from 4 mice) of naïve control and dopamine-depleted mice. Unpaired $t$-test, $t(52)$ = 2.244, *$p$ = 0.0291 in **g**, and $t(10)$ = 2.418, *$p$ = 0.0362 in **h**. All data presented as mean ± SEM.

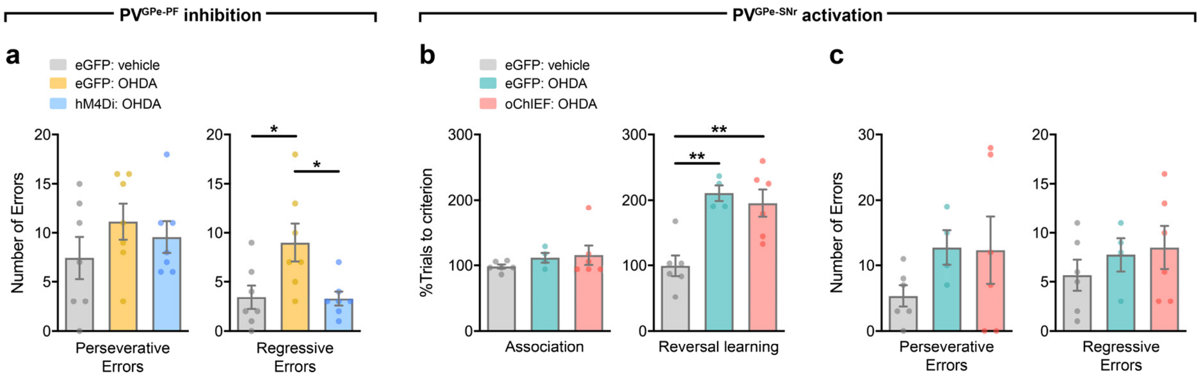


**Extended Data Fig. 9 | Manipulation of PV$^{GPe-PF}$ but not PV$^{GPe-SNr}$ neurons rescues behavioral flexibility deficit in dopamine-depleted mice.** **a**, Number of errors during the reversal-learning phase made by mice that received chemogenetic inhibition in PV$^{GPe-PF}$ neurons after dopamine depletion ($n$ = 7 mice for eGFP-vehicle, $n$ = 7 mice for eGFP-OHDA, and $n$ = 7 mice for hM4Di-OHDA). Left, perseverative errors; One-way ANOVA, $F(2,18) = 0.9771$, $p$ = 0.3955. Right, regressive errors; One-way ANOVA, $F(2,18) = 5.595$, $p = 0.0129$; Bonferroni's *post hoc* test, *$p$ = 0.0312 (eGFP-vehicle vs. eGFP-OHDA) and 0.0267 (eGFP-OHDA vs. hM4Di-OHDA). **b**, Performance of mice that received photostimulation in PV$^{GPe-SNr}$ neurons after dopamine depletion ($n$ = 6 mice for eGFP-vehicle, $n$ = 4 mice for eGFP-OHDA, and $n$ = 6 mice for oChIEF-OHDA). Left, dopamine depletion did not affect performance in the association phase. One-way ANOVA, $F(2,13) = 0.8222$, $p = 0.4611$. Right, activation of PV$^{GPe-SNr}$ neurons during reversal learning did not improved behavioral flexibility in dopamine-depleted mice. One-way ANOVA, $F(2,13) = 11.69$, $p = 0.0012$; Bonferroni's *post hoc* test, **$p$ = 0.0032 (eGFP-vehicle vs. eGFP-OHDA) and 0.0042 (eGFP-vehicle vs. oChIEF-OHDA). **c**, Number of errors during the reversal-learning phase made by mice in **c**. Left, perseverative errors; One-way ANOVA, $F(2,13)$ = 1.308, $p = 0.3038$. Right, regressive errors; One-way ANOVA, $F(2,13) = 0.6464$, $p = 0.54$. All data presented as mean ± SEM.
